# Multi-animal pose estimation and tracking with DeepLabCut

**DOI:** 10.1101/2021.04.30.442096

**Authors:** Jessy Lauer, Mu Zhou, Shaokai Ye, William Menegas, Tanmay Nath, Mohammed Mostafizur Rahman, Valentina Di Santo, Daniel Soberanes, Guoping Feng, Venkatesh N. Murthy, George Lauder, Catherine Dulac, Mackenzie W. Mathis, Alexander Mathis

## Abstract

Estimating the pose of multiple animals is a challenging computer vision problem: frequent interactions cause occlusions and complicate the association of detected keypoints to the correct individuals, as well as having extremely similar looking animals that interact more closely than in typical multi-human scenarios. To take up this challenge, we build on DeepLabCut, a popular open source pose estimation toolbox, and provide high-performance animal assembly and tracking—features required for robust multi-animal scenarios. Furthermore, we integrate the ability to predict an animal’s identity directly to assist tracking (in case of occlusions). We illustrate the power of this framework with four datasets varying in complexity, which we release to serve as a benchmark for future algorithm development.

## Introduction

Advances in sensor and transmitter technology, data mining, and computational analysis herald a golden age of animal tracking across the globe (1). Computer vision is a crucial tool for identifying, counting, as well as annotating animal behavior (2–4). For the computational analysis of fine-grained behavior, pose estimation is often a crucial step, and deep-learning based tools have quickly impacted neuroscience, ethology, and medicine (5, 6).

Many experiments in biology—from parenting mice to fish schooling—require measuring interactions among multiple individuals. Multi-animal pose estimation raises several challenges that can leverage advances in machine vision research, and yet others that need new solutions. In general, the process requires three steps: pose estimation (i.e., keypoint estimation, which is typically done frame-by-frame), assemble (or localize) the individual animal, and then track them through frames. Firstly, due to the interactions of animals there will be occlusions. To make the feature detectors (i.e., the pose estimation step) robust to these altered scene statistics, one can annotate frames with interacting animals. Secondly, one needs to associate detected keypoints to particular individuals. Here, many solutions have been proposed, such as part affinity fields (7), associative embeddings (8, 9), transformers (10) and other mechanisms (11, 12). These are called bottom-up approaches, as detections and links are predicted from the image and the individuals are then “assembled” (typically) in a post-processing step. The alternative, called a top-down approach (e.g., 13, 14), is to first detect individual animals and apply standard pose estimation within the identified regions (reviewed in 15). The utility is often limited in scenarios where the individuals interact closely and occlude one another (7, 13), making individual detections hard. Thirdly, corresponding poses between adjacent frames should be consistently identified and tracked—a task made difficult because of appearance similarity, highly non-stationary behaviors, and possible occlusions. Building on human pose estimation research, some recent packages for multi-animal pose estimation have emerged (16–18). Here, we build on the top-performing animal pose networks, introduce new networks, and compare the current state-of-the-art network on COCO (19) to our model on four animal datasets.

In an effort to make a high-performance yet universal tool, we address the multi-animal pose estimation and tracking challenges by building on bottom-up linking of keypoints to an individual for small animal groups (we demonstrate it for up to fourteen). We developed a new framework by expanding DeepLabCut (20, 21), a popular open source toolbox. Our contributions are as follows:

1. Introduce four datasets of varying difficulty for benchmarking multi-animal pose estimation networks.
2. A novel multi-task architecture that predicts multiple conditional random fields and therefore can predict keypoints, limbs, as well as animal identity.
3. A novel data-driven method for animal assembly that finds the optimal skeleton without user input, and that is state-of-the art (compared to top models on COCO).
4. A new tracking module that is locally and globally optimizable.
5. We show that one can predict the identity of animals, which is useful to link animals across time when temporally-based tracking fails.
6. We extend the open source DeepLabCut software to multi-animal scenarios and provide new graphical user interfaces (GUIs) to allow keypoint annotation and check reconstructed tracks.

## Results

Multi-animal pose estimation can be naively be cast as a data assignment problem in the spatial and temporal domains, where one would need to detect keypoints and identify which individual they belong to (spatial), and further link these keypoints temporally across frames. Thus, to tackle the generic multi-animal pose estimation scenario, we designed a practical, almost entirely data-driven solution that breaks down the larger goal into the smaller sub-tasks of: keypoint estimation, animal assembly, local tracking, and global “tracklet” stitching (Figure S1). To benchmark our pipeline, we also made four datasets.

### Four diverse multi-animal datasets

We considered four multi-animal experiments to broadly validate our approach: three mice in an open field, home-cage parenting in mice, pairs of marmosets housed in a large enclosure, and fourteen fish in a flow tank. These datasets encompass a wide spectrum of behaviors, presenting difficult and unique computational challenges to pose estimation and tracking (Figures 1a, S2). The three mice frequently contact and occlude one another. The parenting dataset contained a single animal with unique keypoints in close interaction with two pups hardly distinguishable from the background or the cotton nest, which also leads to occlusions. The marmoset dataset comprises periods of occlusion, close interactions, highly nonstationary behavior, motion blur, and changes in scale. Likewise, the fish school along all dimensions of the tank, hiding each other in very cluttered scenes, and occasionally leaving the camera’s field of view. We annotated from 5 to 15 body parts of interest depending on the dataset (Figure 1a), in multiple frames for cross-validating the pose estimation and assembly performance, as well as semi-automatically annotated several full videos for evaluating the tracking performance (Table 1). Then for each dataset we created a random split with 95% of the data used for training and the rest for testing. We used this split throughout and share the training data as a collective multi-animal benchmark.

**Figure 1.**
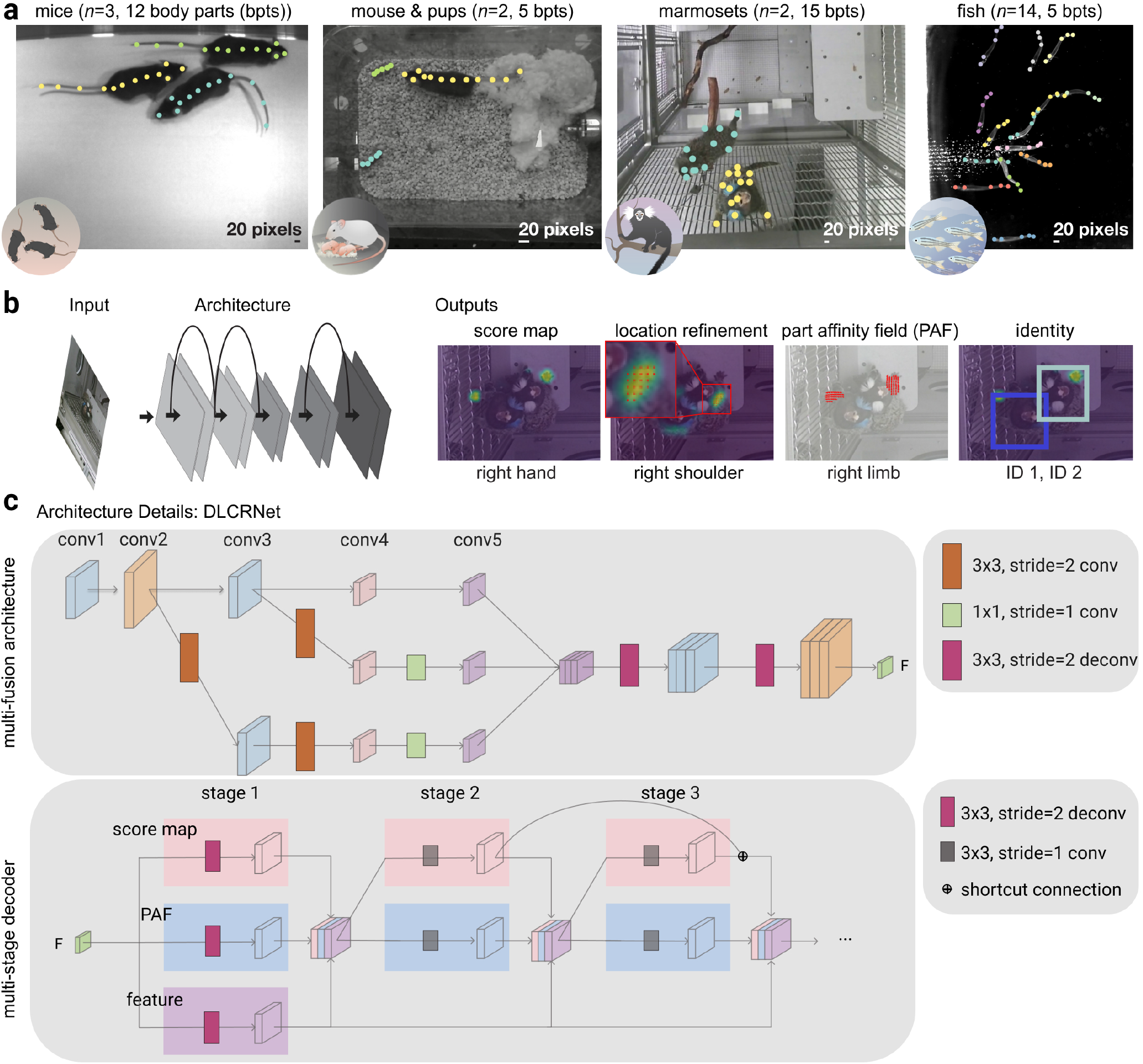
Multi-animal DeepLabCut architecture and benchmarking datasets. **(a)**: Example (cropped) images with (manual) annotations for the four datasets utilized: mice in an open field arena, parenting mice, pairs of marmosets, and schooling fish. **(b)**: A schematic of the general pose estimation module: The architecture is trained to predict the keypoint locations, part affinity fields and animal identity. Three output layers per keypoint predict the probability that a joint is in a particular pixel (score map) as well as shifts in relation to the discretized output map (location refinement field). Furthermore, part affinity fields predict vector fields encoding the orientation of a connection between two keypoints. Example predictions are overlaid on the corresponding (cropped) marmoset frame. The part affinity field for the the right limb helps linking the right hand and shoulder keypoints to the correct individual. **(c)**: The new DeepLabCut architecture contains a multi-fusion module and a multi-stage decoder. In the Multi-fusion module we add the high-resolution representation (conv2, conv3) to low-resolution representation (conv5). The features from conv2 and conv3 are down-sampled by two and one 3×3 convolution layer, respectively to match the resolution of conv5. Before concatenation the features are down-sampled by a 1×1 convolution layer to reduce computational costs and (spatially) up-sampled by two stacked 3×3 deconvolution layers with stride 2. The Multi-stage decoder predicts score maps and part affinity fields (PAF). At the first stage, the feature map from the multi-fusion module are up-sampled by a 3×3 deconvolution layer with stride 2, to get the score map, PAF, the up-sampled feature. In the latter stages, the predictions from the two branches (score maps and PAFs), along with the up-sampled feature are concatenated for the next stage. We applied a shortcut connection between the consecutive stage of the score map. The shown variant of DLCRNet has overall stride 2 (in general, this can be modulated from 2 - 8).

**Table 1.** Multi-animal pose estimation dataset characteristics.

| Feature | Mouse | Pups | Marmoset | Fish |
| --- | --- | --- | --- | --- |
| Labeled frames | 161 | 542 | 7,600 | 100 |
| keypoints | 12 | 5 (+12) | 15 | 5 |
| Individuals | 3 | 2 (+1) | 2 | 14 |
| Identity | no | no | yes | no |
| Ann. video frames | 11645 | 2670 | 15000 | 1100 |
| Total duration (s) | 385 | 180 | 600 | 36 |

Number of labeled training frames, keypoints, and individuals. Animal’s identity was only annotated for the marmosets. The total number of densely human-annotated video frames (and their combined duration in seconds) is also indicated. The annotated videos were used for tracking evaluation.

### Assembling individuals: spatial grouping

#### Multi-task convolutional architectures

We developed multi-task convolutional neural networks (CNN) that perform pose estimation by localizing keypoints in images. This is achieved by predicting *score maps*, which encode the probability that a keypoint occurs at a particular location, as well as *location refinement fields* that predict offsets to mitigate quantization errors due to downsampled score maps (11, 20, 21). Then, in order to group the keypoints to the animal they belong to, we designed the networks to also predict “limbs”, or *part affinity fields*. This task, achieved via additional deconvolution layers, is inspired by OpenPose (7). The intuition behind it is that, in scenarios where multiple animals are present in the scene, learning to predict the location and orientation of limbs will help group pairs of keypoints belonging to an individual. Moreover, we also introduce an output that allows for animal re-identification from visual input directly. This is important in the event of animals that are untrackable using temporal information alone, e.g., when exiting/re-entering the scene (Figure 1b).

Specifically, we adapted ImageNet-pretrained ResNets (22), the current state-of-the art model on the ImageNet benchmark, EfficientNets (23), and introduce a novel multiscale architecture (DLCRNet_ms5) we developed, which is loosely inspired by HRNet (9, 14) for feature extraction (Figure 1c). We then utilize customized multiple parallel deconvolution layers to predict the location of keypoints as well as what keypoints are connected in a given animal (Figure 1b). Ground truth data of annotated keypoints is then used to calculate target score maps, location refinement maps, part affinity fields and to train the network to predict those outputs for a given input image (Figure 1b,c) with augmentation as outlined in the Methods.

#### Keypoint detection & part affinity performance

First, we demonstrate that the architectures perform well for localizing keypoints. We trained independent networks for each dataset and split and evaluated their performance. For each frame and keypoint, we calculated the root-mean-square error between the detections and their closest ground truth neighbors. All the keypoint detectors performed well, with, for example, ResNet-50 having 90% of the prediction errors under 5.1, 10.0, 11.9, and 5.8 pixels for the tri-mouse, parenting, marmoset and fish datasets, respectively (Figure 2a; the scale of these data are shown in Figure 1a). DeepLabCut’s EfficientNet backbones and our new architecture, DL-CRNet_ms5, grant on average a further ~ 21% and ~ 22% reduction in RMSE, respectively (Figure S3a). To ease interpretation, errors were also normalized to 33% of the tip–gill distance for the fish dataset, and 33% of the left-to-right ear distance for the remaining ones (see Methods). We found that 97.0 ± 3.6% of the predictions on the test images were within those ranges (percentage of correct keypoints, PCK; PCK per keypoint are shown in Figure 2a). Thus, DeepLabCut performs well at localizing keypoints in complex, social interactions.

**Figure 2.**
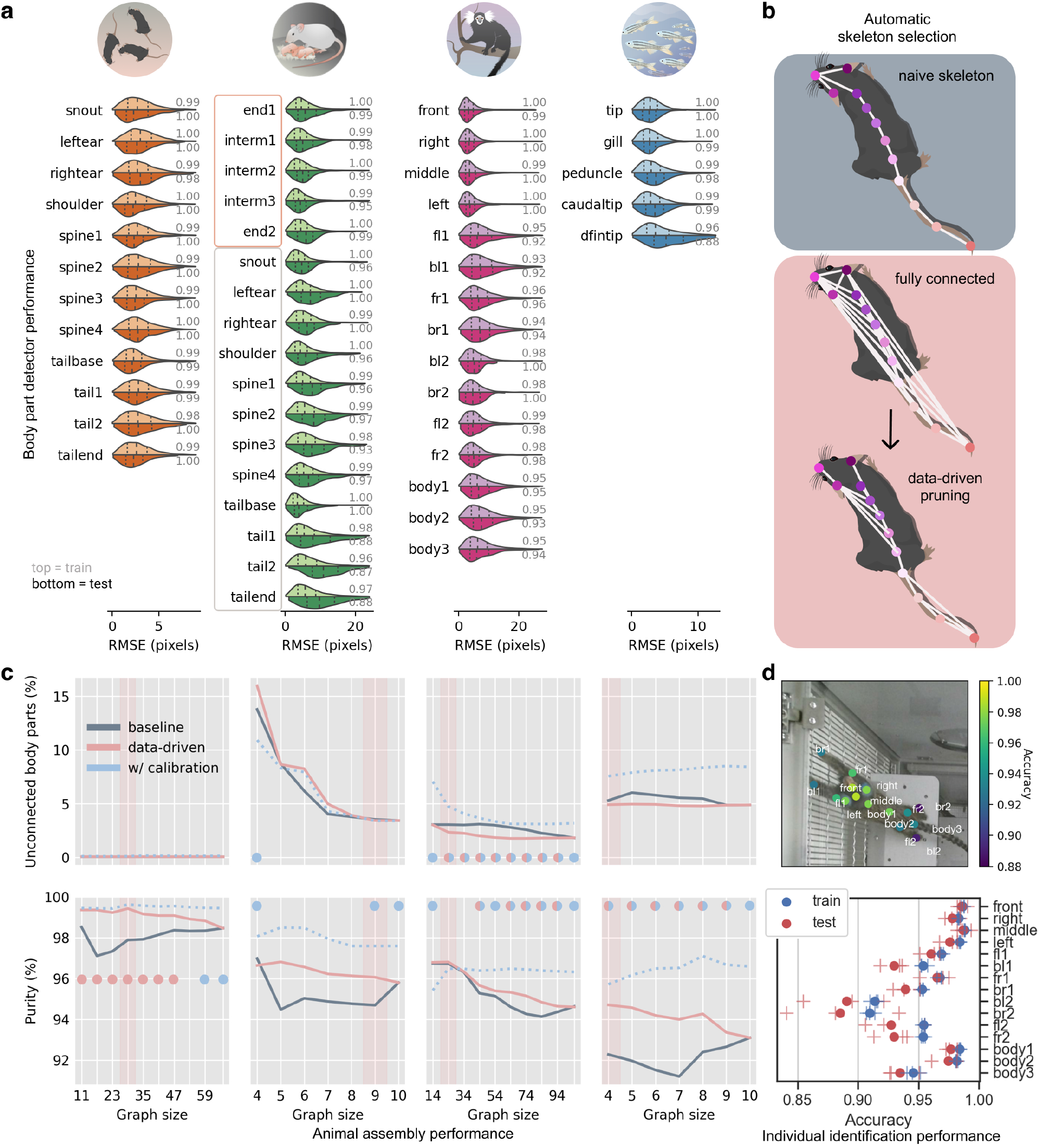
Multi-animal DeepLabCut detector and assembly performance. **(a)**: Distribution of keypoint prediction error for ResNet50_stride8. Violin plots are split vertically in train (top) and test (bottom) errors. Gray numbers indicate PCK. Note that only the first five keypoints of the parenting dataset belong to the pups; the 12 others are keypoints unique to the adult mouse. **(b)**: Illustration of our data-driven skeleton selection algorithm for the mouse skeleton prior to and after automatic pruning. The optimal, minimal skeleton (11 edges; see c) does not resemble a structure one would intuitively draw, and uses some central nodes (e.g., spine2). Mouse from scidraw.io. **(c)**: Animal assembly quality as a function of part affinity graph size for baseline (user-defined) vs data-driven skeleton definitions for multi-stage architecture (DLCRNet_ms5). The top row displays the fraction of keypoints left unconnected after assembly, whereas the bottom row designates the accuracy of their grouping into distinct animals. The colored dots mark statistically significant interactions between graph size and assembling methods, as identified via two-way, repeated-measures ANOVAs. Red dots indicate a significant difference between baseline and data-driven, and blue dots, between data-driven and calibrated assemblies. Light red vertical bars highlight the graph automatically selected to balance the number of body parts left out after assembly and assembly purity. **(d)**: Example test image together with overlaid animal identity prediction accuracy per keypoint averaged over all test images and test splits. With ResNet50_stride8, accuracy peaks at 98.6% for keypoints near the head and drops to 88.5% for more distal parts.

After detection, keypoints need to be assigned to individuals. Thus, we evaluated if the learned part affinity fields helped decide whether two body parts belong to the same or different animals. For example, there are 66 different ways to connect the 12 mouse body parts and many provide high discriminability (Figure S4). We indeed found that predicted limbs were powerful at distinguishing a pair of keypoints belonging to an animal from other (incorrect) pairs linking different mice, as measured by a high auROC (Area Under the Receiver Operating Characteristics) score (0.96 ± 0.04).

#### Data-driven individual assembly performance

Any limb-based assembly approach requires a “skeleton”, i.e., a list of keypoint connections that allows the algorithm to computationally infer which body parts belong together. Naturally, there has to be a path within this skeleton connecting any two body parts, otherwise the body parts cannot be grouped into one animal. Yet, skeletons with additional redundant connections might increase the assembly performance, which raises the question: given the combinatorical nature of skeletons, how should they be picked?^1^ We therefore sought to circumvent the need for arbitrary, hand-crafted skeletons with a method that is agnostic to an animal’s morphology and does not require any user input.

To determine the optimal skeleton, we devised an entirely data-driven method. A network is first trained to predict all graph edges and the least discriminative edges are pruned to determine the skeleton (see Methods). We found that this approach yields perhaps non-intuitive skeletons (Figure 2b), but importantly it improves performance. Our data-driven method (with DLCRNet_ms5) outperforms the naive (baseline) method, which also enhances “purity” of the assembly (Table S1) and reduces the number of missing keypoints (Table S2). Comparisons revealed significantly higher assembly purity with automatic skeleton pruning vs naive skeleton definition at most graph sizes, with respective gains of up to 2.2, 0.5, and 2.4 percentage points in the tri-mouse (graph size=17, *p* < 0.001), marmosets (graph size=74, *p* = 0.002), and fish datasets (graph size=4, *p* < 0.001) (Figure 2b,c). We also found our multi-scale architecture (DLCRNet_ms5) gave us an additional boost in mean average precision (mAP) performance (Tables S3, S4, S5, S6).

To accommodate diverse body plans and annotated keypoints for different animals and experiments, our inference algorithm works for arbitrary graphs. Furthermore, animal assembly achieves at least ≈ 400 frames per second in scenes with fourteen animals, and up to 2000 for small skeletons in 2 or 3 animals (Figure S5).

To additionally benchmark our contributions, we compared our methods to current state-of-the-art methods on COCO (19), a challenging, large-scale multi-human pose estimation benchmark. Specifically we considered HRNet-AE as well as ResNet-AE (see Methods). Importantly, our models performed better than these state-of-the-art methods (Figure S6).

#### Predicting animal identity from images

Animals sometimes differ visually; e.g., due to distinct coat patterns, because they are marked, or carry different instruments (such as an integrated microscope (24)). To allow DeepLabCut to take advantage of such scenarios and improve tracking later on, we developed a head that learns the identity of animals with the same CNN. To benchmark the ID output, we focused on the marmoset data, where (for each pair) one marmoset had light blue dye applied to its tufts. ID prediction accuracy on the test images ranged from > 0.98 for the keypoints closest to the marmoset’s head and its marked features to 0.89 for more distal keypoints. Different backbones can further improve identification performance. While EfficientNet-B0 offers performance comparable to ResNets (~ 0.96), EfficientNet-B7 performs at an average accuracy of 0.99 and 0.98 on the train and test images, respectively (Figure S3b).

### Tracking of individuals: temporal grouping

Once keypoints are assembled into individual animals, the next step is to link them temporally. In order to measure performance in the next steps, entire videos (1 from each dataset) were manually refined to form ground truth sequences, which allowed for the evaluation of tracking and stitching performance ((Figure 3a, and Table 1). Reasoning over the whole video for tracking individuals is not only extremely costly, but also unnecessary. For instance, when animals are far apart, it is straightforward to link each one correctly across time. Thus, we devised a divide-and-conquer strategy. We utilize a simple, online tracking approach to form reliable “tracklets” from detected animals in adjacent frames. Difficult cases (e.g., when animals are closely interacting or after occlusion) often interrupt the tracklets, causing ambiguous fragments that cannot be easily temporally linked. We address this crucial issue post-hoc by optimally stitching tracklets using multiple spatio-temporal cues.

**Figure 3.**
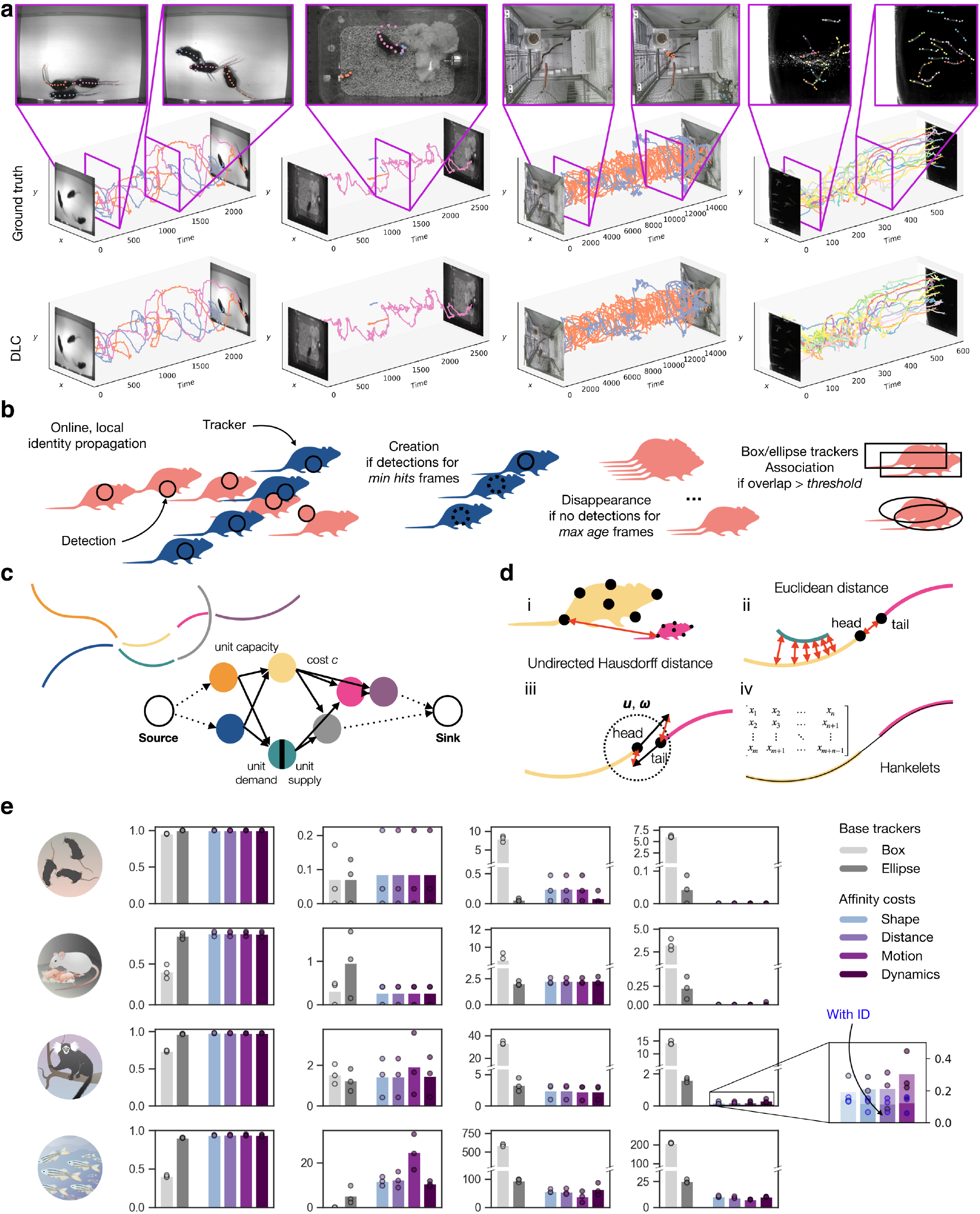
Tracking multiple animals with DeepLabCut. **(a)**: Ground truth and reconstructed animal tracks, together with video frames illustrating representative scene challenges. **(b)**: The identities of animals detected in a frame are propagated across frames using local matching between detections and trackers. **(c)**: Tracklets are represented as nodes of a graph, whose edges encode the likelihood that the connected pair of tracklet belongs to the same track. **(d)**: Four cost functions modeling the affinity between tracklets are implemented: (i) shape similarity using the undirected Hausdorff distance between finite sets of keypoints; (ii) spatial proximity in Euclidean space; (iii) motion affinity using bidirectional prediction of a tracklet’s location; and (iv) dynamic similarity via Hankelets and time-delay embedding of a tracklet’s centroid. **(e)**: Tracklet stitching performance vs box and ellipse tracker baselines, using Multi-Object Tracking Accuracy (MOTA), as well as rates of false positive (FP), false negative (FN) and identity switch expressed in count per 100 frames. Inset: Incorporating identity prediction in the stitching task further reduces the number of switches and improves full track reconstruction. Total number of frames: tri-mouse, 2,330; parenting, 2,670; marmosets, 15,000; fish, 601.

#### Local animal tracking to create tracklets

Assembled animals are linked across frames to form tracklets, i.e., fragments of full trajectories. This task entails the propagation of an animal’s identity in time by finding the optimal association between an animal and its predicted location in the adjacent frame (Figure 3b). The prediction is made by a “tracker”; a lightweight estimator modeling an animal’s state, such as its displacement and velocity. In particular, we implemented a box and an ellipse tracker (see Methods). Whereas the former is standard in object tracking literature (e.g., (25, 26)), we recognized the sensitivity of its formulation to outlier detections (as it is mostly used for pedestrian tracking). Thus, the ellipse tracker was introduced to provide a more robust solution as well as a finer parametrization of an animal’s geometry. Differences in their performance is striking: the ellipse tracker behaves systematically better, reaching near perfect multi-object tracking accuracy and a ~ 8× lower false negative rate, while producing on average ~ 9 × less identity switches (Figure 3e).

#### Globally optimal tracking: tracklet stitching

Because of occlusions, dissimilarity between an animal and its predicted state, or other challenging yet common multianimal tracking issues, tracklets can be interrupted and therefore rarely form complete tracks. The remaining challenge is to stitch these sparse tracklets so as to guarantee continuity and kinematic consistency. Our novel approach is to cast this task as a global minimization problem, where connecting two candidate tracklets incurs a cost inversely proportional to the likelihood that they belong to the same track. Advantageously, the problem can now be elegantly solved using optimization techniques on graph and affinity models (Figure 3c,d).

Compared to only local tracking, we find that our stitching method successfully solves all switches in the tri-mouse and parenting datasets, and reduces them by a factor of ~ 3–9 down to 9 and 0.2 switches/100 frames for the very challenging fish and marmosets datasets, respectively (Figure 3e). To handle a wide range of scenarios, multiple cost functions were devised to model the affinity between a pair of tracklets on the basis of their shape, proximity, motion, and/or dynamics. Furthermore, incorporating visual identity information predicted from the CNN further halved the number of switches (Figure 3e). Example videos with predictions are shown (Supplementary Videos).

### DeepLabCut workflow and usability

We have detailed various new algorithms for solving multianimal pose estimation. Those tools are available in the DeepLabCut GitHub repository and the general workflow was expanded to accommodate multi-animal pose estimation projects for labeling, refining tracklets etc. (Figure S1). The work presented in this paper is termed “maDeepLabCut” and is integrated into version 2.2 code base at [GitHub](https://github.com/DeepLabCut/DeepLabCut) and the Python Package Index (PyPi). We provide Google Colab Notebooks, full project management software and graphic user interface(s), and tooling to run this workflow on cloud computing resources. Moreover, in the code we provide 3D support for multi-animal pose estimation (via multi-camera use), plus this multi-animal variant can be integrated with our real-time software, DeepLabCut-Live! (27). Namely, as we have shown, assembly is fast (Figure S5) and the (local) tracking algorithm we used is an online method, which should allow for real-time experiments.

## Discussion

Here we introduced a multi-animal pose estimation and tracking system by extending DeepLabCut (20, 21, 28) and by building on advances in computer vision, in particular OpenPose (7, 29), EfficientNet (23), HRNet (9, 14), and SORT (25). Firstly, we developed more powerful CNNs (DL-CRNet_ms5), that are state-of-the-art in animal pose and assembly. Secondly, due to the highly variable body shapes of animals (and different keypoints that users might annotate), we developed a novel, data-driven way to automatically find the best skeleton for animal assembly. Thirdly, we proposed fast trackers that (unlike SORT) also reason over long time scales and are more robust to the body plan. Thereby our framework integrates various costs related to movement statistics, and the learned animal identity. We showed that the expanded DeepLabCut toolbox works well for tracking and pose estimation across multiple applications from parenting mice to schools of fish. We also release these datasets (which we have shown to vary in challenges) as benchmarks for the larger community. Our method is flexible and cannot only deal with multiple animals (with one body plan), but also with one agent dealing with multiple others (as in the case of the parenting mouse).

While the computational complexity of our bottom-up approach could limit speed in presence of a large number of animals, we have found it to be on average greater than 400 FPS even with 14 animals. If insufficient, one could resort to top-down approaches (although this tends to work better for videos with few occlusions). In such cases, trackers as idtracker.ai (30), TRex (31) or an object detection algorithm (32) could ideally be used to create bounding boxes around animals prior to estimating poses on these cropped images as already possible with “vanilla” DeepLabCut, DeepPoseKit (33), etc. (discussed in Mathis et al. (15) and Walter and Couzin (31)).

In summary, we report the development and performance of a new multi-animal animal pose estimation pipeline. We integrated and developed state-of-the-art neural network architectures, developed a novel data-driven pipeline that not only optimizes performance, but also does not require extensive domain knowledge. Lastly, with the 4 datasets we release here (> 8,000 labeled frames), we also aim to help advance the field of animal pose estimation in the future.

## Acknowledgments

Funding was primarily provided by the Rowland Institute at Harvard University (MWM, TN, AM, JL), the Chan Zuckerberg Initiative DAF (MWM, AM, JL), and EPFL (MWM, AM). Dataset collection was funded by: Office of Naval Research grants N000141410533 and N00014-15-1-2234 (GVL), HHMI and NIH grant 2R01HD082131 (MMR, CD); NIH grant 1R01NS116593-01 (MMR, CD and VNM) We are grateful to Maxime Vidal for converting datasets. We thank the beta testers and DLC community for feedback and testing. MWM is the Bertarelli Foundation Chair of Integrative Neuroscience.

## Author contributions

Conceptualization (AM, MWM), Formal analysis and code (JL, AM), novel deep architectures (MZ, SY, AM), GUIs (JL, MWM, TN), Marmoset data (WM, GF), Parenting data (MMR, AM, CD), Tri-mouse data (DS, AM, VNM), Fish data (VD, GL), Writing (AM, JL, MWM) with input from all authors.

## Methods

### Multi-animal datasets

For this study we established four differently challenging multi-animal datasets from ecology and neuroscience.

#### Tri-mouse dataset

Three wild-type (C57BL/6J) male mice ran on a paper spool following odor trails (20). These experiments were carried out in the laboratory of Venkatesh N. Murthy at Harvard University. Data were recorded at 30Hz with 640 × 480 pixels resolution acquired with a Point Grey Firefly FMVU-03MTM-CS. One human annotator was instructed to localize the 12 keypoints (snout, left ear, right ear, shoulder, four spine points, tail base and three tail points). To have smaller frames (for training with larger batch sizes), and more diverse dataset each image was used to randomly create 10 images of size 400 × 400, which are picked as subsets of the original image—this can be done automatically (using the utility function `deeplabcut.cropimagesandlabels`).

#### Parenting behavior

Parenting behavior is a pup directed behavior observed in adult mice involving complex motor actions directed towards the benefit of the offspring (34, 35). These experiments were carried out in the laboratory of Catherine Dulac at Harvard University. The behavioral assay was performed in the homecage of singly housed adult female mice in dark/red light conditions. For these videos, the adult mice was monitored for several minutes in the cage followed by the introduction of pup (4 days old) in one corner of the cage. The behavior of the adult and pup was monitored for a duration of 15 minutes. Video was recorded at 30Hz using a Microsoft LifeCam camera (Part #: 6CH-00001) with a resolution of 1280 × 720 pixels or a Geovision camera (model no.: GV-BX4700-3V) also acquired at 30 frames per second at a resolution of 704 × 480 pixels. A human annotator labeled on the adult animal the same 12 body points as in the tri-mouse dataset, and five body points on the pup along its spine. Initially only the two ends were labeled, and intermediate points were added by interpolation and their positions was manually adjusted if necessary. Similar to the tri-mouse dataset, we created random crops of 400 × 400 pixels before training. All surgical and experimental procedures for mice were in accordance with the National Institutes of Health Guide for the Care and Use of Laboratory Animals and approved by the Harvard Institutional Animal Care and Use Committee.

#### Marmoset home-cage behavior

All animal procedures are overseen by veterinary staff of the MIT and Broad Institute Department of Comparative Medicine, in compliance with the NIH guide for the care and use of laboratory animals and approved by the MIT and Broad Institute animal care and use committees. Video of common marmosets (*Callithrix jacchus*) was collected in the laboratory of Guoping Feng at MIT. Marmosets were recorded using Kinect V2 cameras (Microsoft) with a resolution of 1080p and frame rate of 30 Hz. After acquisition, images to be used for training the network were manually cropped to 1000 × 1000 pixels or smaller. For our analysis, we used 7,600 labeled frames from 40 different marmosets collected from 3 different colonies (in different facilities). Each cage contains a pair of marmosets, where one marmoset had light blue dye applied to its tufts. One human annotator labeled the 15 marker points on each animal present in the frame (frames contained either 1 or 2 animals).

#### Fish schooling behavior

Schools of inland silversides (*Menidia beryllina*, n=14 individuals per school) were recorded in the Lauder Lab at Harvard University while swimming at 15 speeds (0.5 to 8 BL/s, body length, at 0.5 BL/s intervals) in a flow tank with a total working section of 28 × 28 × 40 cm as described in previous work (36), at a constant temperature (18±1°C) and salinity (33 ppt), at a Reynolds number of approximately 10,000 (based on BL). Dorsal views of steady swimming across these speeds were recorded by high-speed video cameras (FASTCAM Mini AX50, Photron USA, San Diego, CA, USA) at 60-125 frames per second (feeding videos at 60 fps, swimming alone 125 fps). The dorsal view was recorded above the swim tunnel and a floating Plexiglas panel at the water surface prevented surface ripples from interfering with dorsal view videos. Random crops of 400 × 400 pixels were created. Five keypoints were labeled (tip, gill, peduncle, dorsal fin tip, caudal tip).

#### Dataset properties

All frames were labeled with the annotation GUI; depending on the dataset between 100 and 7,600 frames were labeled (Table 1). We illustrated the diversity of the postures by clustering (Figure S2). To assess the level of interactions, we evaluate a Proximity Index (S2m), whose idea is inspired from (13) but its computation was adapted to keypoints. For each individual, instead of delineating bounding boxes to determine the vicinity of an animal we rather define a circle centered on the individual’s centroid and of sufficiently large radius such that all of that individual’s keypoints are inscribed within the circle; this is a less static description of the immediate space an animal can reach. The index is then taken as the ratio between the number of keypoints within that region that belong to other individuals and the number of keypoints of the individual of interest (Figure S2m).

For each dataset we created three random splits with 95% of the data used for training and the rest for testing. The first one was used throughout and will be made available as a benchmark. Note that identity prediction accuracy (2d) and tracking performance (3e) are reported on all three splits, and all show little variability.

#### Pose estimation

##### Multi-task deep learning architecture

DeepLabCut consists of keypoint detectors, comprising a deep convolutional neural network (CNN) pretrained on ImageNet as a backbone together with multiple deconvolutional layers (11, 20, 28). Here, as backbones we considered Residual Networks (ResNet) (22), and EfficientNets (23, 28). Other backbones are integrated in the toolbox (28) such as MobileNetV2 (37). We utilize a stride of 16 for the ResNets (achieved by atrous convolution) and then upsample the filter banks by a factor of two to predict the score maps and location refinement fields with an overall stride of 8. Furthermore, we developed a multi-scale architecture that upsamples from conv5 and fuses those filters with filters learned as 1 × 1 convolutions from conv3. This bank is then upsampled by a factor of 2 via deconvolution layers. This architecture thus learns from multiple scales with an overall stride of 4 (including the up-sampling in the decoder). We implemented a similar architecture for EfficientNets. These architectures are called ResNet50_strideX and (EfficientNet) bY_strideX for strides 4 and 8; we used ResNet50 and B0 and B7 for experiments (Figure S3).

We further developed a multi-scale architecture (DLCR-Net_ms5) which fuses high resolution feature map to lower resolution feature map (Figure 1c)—we concatenated the feature map from conv5, the feature map learned as a 3× 3 convolutions followed by a 1 × 1 convolutions from conv3 and the feature map learned as 2 stacked 3× 3 convolutions and a 1 × 1 convolutions from conv2. This bank is then upsampled via (up to) 2 deconvolution layers. Depending on how many deconvolution layers are used this architecture learns from multiple scales with an overall stride of 2-8 (including the up-sampling in the decoder). For most cases we found significant improvements with this architecture typically for stride 4 (see Results).

DeepLabCut creates three output layers per keypoint that encode an intensity and a vector field. The purpose of the deconvolution layers is to upsample the spatial information (Figure 1b,c). Consider an input image *I*(*x, y*) with ground truth keypoint (*x^k^, y^k^*) for index *k*. One of the output layers encodes the confidence of a keypoint *k* being in a particular location (*S^k^*(*p, q*)), and the other two layers encode the (x-) and (y-) difference (in pixels of the full-sized) image between the original location and the corresponding location in the downsampled (by the overall stride) location 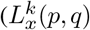 and 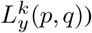. For each training image the architecture is trained end-to-end to predict those outputs. Thereby, the ground truth keypoint is mapped into a target score map, which is one for pixels closer to the target (this can be subpixel location) than radius *r* and 0 otherwise. We minimize the cross entropy loss for the score map (*S^k^*) and the location refinement loss calculated Huber loss (11, 20).

To link specific keypoints within one animal, we employ part affinity fields (PAF), which were proposed by Cao et al. (7). Each (ground truth) PAF 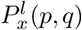 and 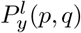 for limb/connection *l* connecting keypoint *k_i_* and *k_j_* places a directional unit vector at every pixel vector within a predefined distance from the ideal line connecting two keypoints (modulated by `pafwidth`). We trained DeepLabCut to also minimize the *L*1-loss between the predicted and true PAF, which is added to the other losses.

Inspired by Cao et al. (7), we refine the score maps and PAFs in multiple stages. As can be seen from Figure 1b, at the first stage, the original image feature from the backbone are fed into the network to predict the score map, PAF and the feature map. The output of each branch, concatenated with the feature map is fed into the subsequent stages. However, unlike Cao et al., we observed that simply adding more stages can cause performance degradation. To overcome that, we introduced shortcut connections between two consequence stages on the score map branch to improve multiple stage prediction.

Examples for score maps, location refinement and PAFs are shown in Figure 1b. For training, we used the Adam optimizer (38) with batch size 4 and learning schedule (0.0001 for first 7,500 iterations then 5*e* – 05 until 12,000 iterations and then 1*e* – 05) unless otherwise noted. We trained for 60,000 (batch size 8); this was enough to reach good performance (Figures 2a and S3). During training we also augmented images by using techniques including cropping, rotation, covering with random boxes, and motion blur.

##### CNN-based identity prediction

For animal identification we used a classification approach (4), while also considering spatial information. To have a monolithic solution (with just a single CNN), we simply predict in parallel the identity of each animal from the image. For this purpose, *n* deconvolution layers are added for *n* individuals. The network is trained to predict the summed score map for all keypoints of that individual. At test time, we then look up which of the output classes has the highest likelihood (for a given keypoint) and assign that identity to the keypoint. This output is trained jointly in a multi-task configuration. We evaluate the performance for identity prediction on the marmoset dataset (Figure 2d).

##### Multi-animal inference

Any number of keypoints can be defined and labeled with the toolbox; additional ones can be added later on. We recommend labeling more keypoints than a subsequent analysis might require, since it improves the part detectors (20) and, more importantly, animal assembly as seen below.

For each keypoint one obtains the most likely keypoint location (*x*,y\**) by taking the maximum: (*p*,q\**) = argmax_(*p,q*_)*S^k^*(*p,q*) and computing:

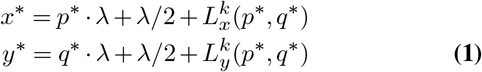

with overall stride λ. If there are multiple keypoints *k* present then one can naturally take the local maxima of *S^k^* to obtain the corresponding detections.

Thus, one obtains putative keypoint proposals from the score maps and location refinement fields. We then use the part affinity fields to assign the cost for linking two keypoints (within a putative animal). For any pair of keypoint proposals (that are connected via a limb as defined by the part affinity graph) we evaluate the affinity cost by integrating along line *γ* connecting two proposals, normalized by the length of *γ*:

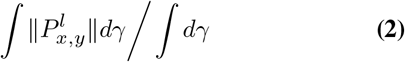

This integral is computed by sampling. Thus, for a given part affinity graph, one gets a (possibly) large number of detections and costs. The next step is to assemble those detections into animals.

##### Data-driven part affinity field graph selection

To relieve the user from manually defining connections between keypoints, we developed an entirely data-driven procedure. Models are trained on a complete graph in order to learn all possible body part connections. The graph is then pruned based on edge discriminability power on the training set. For this purpose, within- and between-animal part affinity cost distributions (bin width=0.01) are evaluated (see Figure S4 for the mouse dataset). Edges are then ranked in decreasing order of their ability to separate both distributions—evaluated from the area under the ROC curve. The smallest, data-driven graph is taken as the maximum spanning tree (i.e., a subgraph covering all keypoints with the minimum possible number of edges that also maximizes part affinity costs). For graph search following a network’s evaluation, up to nine increasingly redundant graphs are formed by extending the minimal skeleton progressively with strongly discriminating edges in the order determined above. By contrast, baseline graphs are grown from a skeleton a user would naively draw, with edges iteratively added in reversed order (i.e., from least to most discriminative). The graph jointly maximizing purity and the fraction of connected keypoints is the one retained to carry out the animal assemblies.

##### Animal assembly

Animal assembly refers to the problem of assigning keypoints to individuals. Yet, reconstructing the full pose of multiple individuals from a set of detections is NP hard, as it amounts to solving a *k*-dimensional matching problem (a generalization of bipartite matching from 2 to *k* disjoint subsets) (7, 39). To make the task more tractable, we break the problem down into smaller matching tasks, in a manner akin to Cao et al. (7).

For each edge type in the data-driven graph defined earlier, we first pick strong connections based on affinity costs alone. Following the identification of all optimal pairs of keypoints, we seek unambiguous individuals by searching this set of pairs for connected components—in graph theory, these are subsets of keypoints all reachable from one another but that do not share connection with any additional keypoint; consequently, only connectivity, but not spatial information, is taken into account. Breadth-first search runs in linear time complexity, which thus allows the rapid pre-determination of unique individuals. Note that, unlike (7), redundant connections are seamlessly handled and do not require changes in the formulation of the animal assembly.

Then, remaining connections are sorted in descending order of their affinity costs (Eqn2) and greedily linked. To further improve the assembly’s robustness to ambiguous connections (that is, a connection attempting to either link keypoints belonging to two distinct individuals or overwrite existing ones), the assembly procedure can be calibrated by determining the prior probability of an animal’s pose as a multivariate normal distribution over the distances between all pairs of keypoints. Mean and covariance are estimated from the labeled data via density estimation with Gaussian kernel and bandwidth automatically chosen according to Scott’s Rule. A skeleton is then only grown if the candidate connection reduces the Mahalanobis distance between the resulting configuration and the prior (referred to as w/ calibration in Figure 2c). Lastly, our assembly’s implementation is fully parallelized to benefit greatly from multiple processors (Figure S5).

Optionally (and only when analyzing videos), affinity costs between body parts can be weighted so as to prioritize strong connections that were preferentially selected in the past frames. To this end, and inspired by (40), we compute a temporal coherence cost as follows: 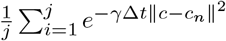 where *γ* controls the influence of distant frames (and is set to 0.01 by default); *c* and *c_n_* are the current connection and its closest neighbor in the relevant past frame; and Δ*t* is the temporal gap separating these frames.

##### Detection performance and evaluation

To compare the human annotations with the model predictions we used the Euclidean distance to the closest predicted keypoint (root mean square error, abbreviated: RMSE) calculated per keypoint. Depending on the context this metric is either shown for a specific keypoint, averaged over all keypoints, or averaged over a set of train/test images (Figures 2a and S3). Nonetheless, unnormalized pixel errors may be difficult to interpret in certain scenarios; e.g., marmosets dramatically vary in size as they leap from the top to the bottom of the cage. Thus, we also calculated the percentage of correct keypoints (PCK) metric (28, 41); i.e., the fraction of predicted keypoints that fall within a threshold distance from the location of the ground truth detection. PCK was computed in relation to a third of the tip–gill distance for the fish dataset, and a third of the left–right ear distance for the remaining ones.

Animal assembly quality was evaluated in terms of mean Average Precision (mAP) computed over object keypoint similarity thresholds ranging from 0.50 to 0.95 in steps of 0.05, as is standard in human pose literature and COCO challenges (19). Keypoint standard deviation was set to 0.1. As interpretable metrics, we also computed the number of ground truth keypoints left unconnected (after assembly) and purity — an additional criterion for quality that can be understood as the accuracy of the assignment of all keypoints of a putative subset to the most frequent ground truth animal identity within that subset (42).

##### Statistics for assessing data-driven method

Two-way, repeated-measures ANOVAs were performed using Pingouin (version 0.3.11 (43)) to test whether graph size and assembling method (naive vs data-driven vs calibrated assembly) had an impact on the fraction of unconnected body parts and assembly purity. Since sphericity was violated, the Greenhouse–Geisser correction was applied. Provided a main effect was found, we conducted multiple post-hoc (paired, two-sided) tests adjusted with Bonferroni correction to locate pairwise differences. The Hedges’ *g* was calculated to report effect sizes between sets of observations.

##### Comparison to state-of-the-art pose estimation models

For benchmarking, we compared our architectures to current state-of-the-art architectures on COCO (19), a challenging, large-scale multi-human pose estimation benchmark. Specifically we considered HRNet (9, 44) as well as ResNet backbones (22) with Associative Embedding (8) as implemented in the well-established MMPose toolbox (45). We chose them as control group for their simplicity (ResNet) and performance (HRNet). We used the bottom-up variants of both model that are implemented in MMPose. The bottom-up variants leverage associative embedding as the grouping algorithms (8). In particular, the bottom-up variant of HRNet we used has mAP that is comparable to the state-of-the-art model HigherHRNet (9) in COCO (69.8 vs. 70.6) for multiple scale test and (65.4 vs. 67.7) for single scale test.

To fairly compare we used the same train/test split. The total training epochs are set such that models from two groups see roughly same number of images. The hyper-parameters search was manually performed to find the optimal hyperparameters. For the small dataset such as tri-mouse and (largest) marmoset, we found that the default settings for excellent performance on COCO gave optimal accuracy except that we needed to modify the total training steps to match DeepLabCut’s. For both the marmoset and tri-mouse datasets, the initial learning rate was 0.0015. For 3 mouse dataset, the total epochs is 3000 epochs and the learning rate decayed by a factor of 10 at at 600 and 1000 epochs. For the Marmoset dataset, we trained for 50 epochs and the learning rate decayed after 20 and 40 epochs. The batch size was 32 and 16 for ResNet-AE and HRNet-AE, respectively. For smaller datasets such as tri-mouse, fish and parenting, we found that a smaller learning rate and a smaller batch size gave better results; a total of 3000 epochs were used. After hyper-parameter search, we set batch size as 4 and initial learning rate a 0.0001, which then decayed at 1000 epochs and 2000 epochs. As within DeepLabCut, multiple scale test and flip test were not performed (which is, however, common for COCO evaluation). For the parenting dataset, MM-Pose models can only be trained on one data set (simultaneously), which is why these models are not trained to predict the mouse, and we only compare the performance on the pups. Full results are shown in Figure S6.

### Animal tracking

Having seen that DeepLabCut provides a strong predictor for individuals and their keypoints, detections are linked across frames using a tracking-by-detection approach (e.g., (46)). Thereby, we follow a divide-and-conquer strategy for (local) tracklet generation and tracklet stitching (Figure S4b,c).

Specifically, we build on the Simple Online and Realtime Tracking framework (SORT; (25)) to generate tracklets. The inter-frame displacement of assembled individuals is estimated via Kalman filter-based trackers. The task of associating these location estimates to the model detections is then formulated as a bipartite graph matching problem solved with the Hungarian algorithm, therefore guaranteeing a one-to-one correspondence across adjacent frames. Note that the trackers are agnostic to the type of skeleton (animal body plan), which render them robust and computationally efficient.

#### Box tracker

Bounding boxes are a common and well-established representation for object tracking. Here they are computed from the keypoint coordinates of each assembled individual, and expanded by a margin optionally set by the user. The state *s* of an individual is parametrized as: 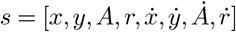, where *x* and *y* are the 2D coordinates of the center of the bounding box; *A*, its area; and *r*, its aspect ratio, together with their first time derivatives. Unlike the original formulation (25), box aspect ratio is allowed to vary over time in order to account for abrupt changes in body shape (e.g., during turns). Association between detected animals and tracker hypotheses is based upon the Intersection-over-Union measure of overlap.

#### Ellipse tracker

A 2*σ* covariance error ellipse is fitted to an individual’s detected keypoints. The state is modeled as: 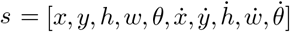, where *x* and *y* are the 2D coordinates of the center of the ellipse; *h* and *w*, the lengths of its semi-axes; and *θ*, its inclination relative to the horizontal. We anticipated that this parametrization would better capture subtle changes in body conformation, most apparent through changes in ellipse width/height and orientation. Moreover, an error ellipse confers robustness to outlier keypoints (e.g., a prediction assigned to the wrong individual, which would cause the erroneous delineation of an animal’s boundary under the above-mentioned Box tracking). In place of the ellipse overlap, the similarity cost *c* between detected and predicted ellipses is efficiently computed as: *c* = 0.8 * (1 – *d*) +0.2 * (1 – *d*) * (cos(*θ_d_* – *θ_p_*)), where *d* is the Euclidean distance separating the ellipse centroids normalized by the length of the longest semi-axis.

The existence of untracked individuals in the scene is signaled by assembled detections with a similarity cost lower than `iou_threshold` (set to 0.6 in our experiments). In other words, the higher the similarity threshold, the more conservative and accurate the frame-by-frame assignment, at the expense of shorter and more numerous tracklets. Upon creation, a tracker is initialized with the required parameters described above, and all (unobservable) velocities are set to 0. To avoid tracking sporadic, spurious detections, a tracker is required to live for a minimum of `min_hits` consecutive frames, or is otherwise deleted. Occlusions and reidentification of individuals are handled with the free parameter `max_age`—the maximal number of consecutive frames tracks can remain undetected before the tracker is considered lost. We set both to 1 to delegate the tasks of tracklet re-identification and false positive filtering to our Tracklet-Stitcher, as we shall see below.

### Tracklet stitching

Greedily linking individuals across frames is locally, but not globally, optimal. An elegant and efficient approach to reconstructing full trajectories (or tracks) from sparse tracklets is to cast the stitching task as a network flow minimization problem (47, 48). Intuitively, each fully reconstructed track is equivalent to finding a flow through the graph from a source to a sink, subject to capacity constraints and whose overall linking cost is minimal (Figure S4c).

#### Formulation

The tracklets collected after animal tracking are denoted as 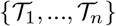, and each contains a (temporally) ordered sequence of observations and time indices. Thereby, the observations are given as vectors of body part coordinates in pixels and likelihoods. Importantly, and in contrast to most approaches described in the literature, the proposed approach requires solely spatial and temporal information natively, while leveraging visual information (e.g., animals’ identities predicted beforehand) is optional (see Figure **??** for marmosets). This way, tracklet stitching is agnostic to the framework poses were estimated with, and works readily on previously collected kinematic data.

We construct a directed acyclic graph *G* = (*V, E*) using NetworkX (49) to describe the affinity between multiple tracklets, where the *i*th node *V_i_* corresponds to the *i*th tracklet 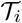, and *E* is the set of edges encoding the cost entailed by linking the two corresponding tracklets (or, in other words, the likelihood that they belong to the same track). In our experiments, tracklets shorter than five frames were flagged as residuals: They do not contribute to the construction of the graph and are incorporated only after stitching. This minimal tracklet length can be changed by a user. To drastically reduce the number of possible associations and make our approach scale efficiently to large videos, edge construction is limited to those tracklets that do not overlap in time (since an animal cannot occupy multiple spatial locations at any one instant) and temporally separated by no more than a certain number of frames. By default, this threshold is automatically taken as 1.5 * *τ*, where *τ* is the smallest temporal gap guaranteeing that all pairs of consecutive tracklets are connected. Alternatively, the maximal gap to consider can be programmatically specified. The source and the sink are two auxiliary nodes that supply and demand an amount of flow *k* equal to the number of tracks to form. Each node is virtually split in half: an input with unit demand and an output with unit supply, connected by a weightless edge. All other edges have unit capacity and a weight *w* calculated from the affinity models described in the next subsection. Altogether, these constraints ensure that all nodes are visited exactly once, which thus amounts to a problem similar to covering *G* with *k* node-disjoint paths at the lowest cost. We considered different affinities for linking tracklets (Figure S4d).

#### Affinity models

##### Motion affinity

Let us consider two non-overlapping tracklets 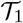 and 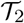 consecutive in time. Their motion affinity is measured from the error between the true locations of their centroids (i.e., unweighted average keypoint) and predictions made from their linear velocities. Specifically, we calculate a tracklet’s tail and head velocities by averaging instantaneous velocities over its three first and last data points (Figure S4d). Assuming uniform, rectilinear motion, the centroid location of 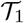 at the starting frame of 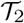 is estimated, and we note *d_f_* the distance between the forward prediction and the actual centroid coordinates. The same procedure is repeated backward in time, predicting the centroid location of 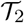 at the last frame of 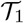 knowing its tail velocity, yielding *d_b_*. Motion affinity is then taken as the average error distance.

##### Spatial proximity

If a pair of tracklets overlaps in time, we calculate the Euclidean distance between their centroids averaged over their overlapping portion. Otherwise, we evaluate the distance between a tracklet’s tail and the other’s head.

##### Shape similarity

Shape similarity between two tracklets is taken as the undirected Hausdorff distance between the two sets of keypoints. Although this measure provides only a crude approximation of the mismatch between two animals’ skeletons, it is defined for finite sets of points of unequal size; e.g., it advantageously allows the comparison of skeletons with a different number of visible keypoints.

##### Dynamic similarity

To further disambiguate tracklets in the rare event that they are spatially and temporally close, and similar in shape, we propose to use motion dynamics in a manner akin to (50). The procedure is fully data-driven, and requires no a priori knowledge of the animals’ behavior. In the absence of noise, the rank of the Hankel matrix—a matrix constructed by stacking delayed measurements of a tracklet’s centroid—theoretically determines the dimension of state space models; i.e., it is a proxy for the complexity of the underlying dynamics (51). Intuitively, if two tracklets originate from the same dynamical system, a single, low-order regressor should suffice to approximate them both. On the other hand, tracklets belonging to different tracks would require a higher-order (i.e., more complex) model to explain their spatial evolution (50). Low rank approximation of a noisy matrix though is a complex problem, as the matrix then tends to be full rank (i.e., all its singular values are nonzero). For computational efficiency, we approximate the rank of a large numbers of potentially long tracklets using singular value decomposition (SVD) via interpolative decomposition. Optimal low rank was chosen as the rank after which eigenvalues drop by less than 1%.

##### Problem solution for stitching

The optimal flow solution can be found using a min-cost flow algorithm. We employ NetworkX’s capacity scaling variant of the successive shortest augmenting path algorithm, which requires polynomial time for the assignment problem (i.e., when all nodes have unit demands and supplies; (52)). Residual tracklets are then greedily added back to the newly stitched tracks at locations that guarantee time continuity and, when there are multiple candidates, minimize the distances to the neighboring tracklets. Note though that residuals are typically very short, making the assignment decisions error-prone. To improve robustness and simultaneously reduce complexity, association hypotheses between temporally close residual tracklets are stored in the form of small directed acyclic graphs during a preliminary forward screening pass. An hypothesis likelihood is then scored based on pairwise tracklet spatial overlap, and weighted longest paths are ultimately kept to locally grow longer, more confident residuals.

This tracklet stitching process is implemented in DeepLabCut and automatically carried out after assembly and tracking. The tracks can then also be manually refined in a dedicated GUI (Figure S1).

##### Tracking performance evaluation

Tracking performance was assessed with the multi-object tracking (MOT) metrics (53). Namely, we retained: multi-object tracking accuracy (MOTA), evaluating a tracker’s overall performance at detecting and tracking individuals (all possible sources of errors considered) independently of its ability to predict an individual’s location; the number of false positives (or false alarms), which signals tracker predictions without corresponding ground truth detections; the number of misses, which counts actual detections for which there are no matching trackers; and, the number of switches (or mismatches), occurring most often when two animals pass very close to one another or if tracking resumes with a different ID after an occlusion.

## Supplemental Materials

Figures and Tables supporting Lauer et al.

**Table S1.** Graph Assembly Statistics: Graph size × assembling method interactions were found to have significant effects on purity in all datasets, with small to medium effect sizes.

| Dataset | F-value | P-value | $\eta_p^2$ |
| --- | --- | --- | --- |
| tri-mouse | 7.170 | <0.001 | 0.043 |
| parenting | 2.895 | 0.038 | 0.019 |
| marmosets | 55.600 | <0.001 | 0.020 |
| fish | 9.584 | <0.001 | 0.088 |

**Table S2.** Fraction of unconnected keypoints: Graph size × assembling method interactions affect the number of unconnected keypoints.

| Dataset | F-value | P-value | $\eta_p^2$ |
| --- | --- | --- | --- |
| tri-mouse | 0.818 | 0.367 | 0.005 |
| parenting | 3.018 | 0.039 | 0.020 |
| marmosets | 90.252 | <0.001 | 0.450 |
| fish | 27.986 | <0.001 | 0.220 |

**Table S3.** Mean Average Precision as a function of graph size for the tri-mouse dataset (more model benchmarking is available in the Suppl. Sheet).

| network name | PW | G | 11 | 17 | 23 | 29 | 35 | 41 | 47 | 53 | 59 | 66 |
| --- | --- | --- | --- | --- | --- | --- | --- | --- | --- | --- | --- | --- |
| DLCRNet MF-3M_s4 | 8 | c | 0.986 | 0.989 | 0.989 | 0.988 | **0.991** | 0.984 | 0.983 | 0.983 | **0.985** | 0.985 |
| DLCRNet MS3-3M_s4 | 16 | c | 0.982 | 0.984 | 0.984 | **0.990** | 0.990 | **0.987** | **0.988** | 0.988 | 0.984 | **0.986** |
| DLCRNet MS3-3M_s2 | 20 | c | 0.949 | 0.971 | 0.969 | 0.988 | 0.987 | 0.978 | 0.984 | **0.990** | 0.960 | 0.961 |
| DLC-ResNet50_s4 | 20 | c | **0.991** | **0.991** | **0.991** | 0.986 | 0.987 | 0.985 | **0.988** | 0.984 | 0.987 | **0.986** |
| DLC-ResNet50_s4 | 20 | d | 0.987 | 0.987 | 0.985 | 0.982 | 0.977 | 0.968 | 0.970 | 0.964 | 0.963 | 0.948 |
| DLC-ResNet50_s8 | 20 | c | 0.956 | 0.979 | 0.979 | 0.982 | 0.981 | 0.981 | 0.984 | 0.984 | 0.978 | 0.979 |
| DLC-EfficientNet-b7_s4 | 20 | c | 0.967 | 0.969 | 0.973 | 0.973 | 0.979 | 0.976 | 0.978 | 0.970 | 0.969 | 0.961 |
| DLC-EfficientNet-b0_s4 | 20 | c | 0.972 | 0.977 | 0.978 | 0.971 | 0.962 | 0.965 | 0.963 | 0.966 | 0.964 | 0.969 |
| DLC-EfficientNet-b0_s8 | 20 | c | 0.968 | 0.971 | 0.972 | 0.974 | 0.975 | 0.976 | 0.970 | 0.969 | 0.968 | 0.969 |

PW = paf width; G = Graph; c = calibrated; d = data-driven; b = baseline; s = stride; MF = multifusion; MS = multistage

**Table S4.** Mean Average Precision as a function of graph size for the parenting dataset (more model benchmarking is available in the Suppl. Sheet.)

| network name | PW | G | 4 | 5 | 6 | 7 | 8 | 9 | 10 |
| --- | --- | --- | --- | --- | --- | --- | --- | --- | --- |
| DLCRNet MS5-s4 | 20 | c | 0.633 | 0.656 | 0.653 | 0.656 | 0.654 | 0.660 | 0.654 |
| DLCRNet MS5-s4 | 20 | b | 0.613 | 0.609 | 0.607 | 0.655 | 0.660 | 0.654 | 0.651 |
| DLCRNet MS5-s4 | 20 | d | 0.604 | 0.609 | 0.647 | 0.650 | 0.660 | 0.655 | 0.654 |
| DLC-ResNet50_s8 | 20 | c | 0.582 | 0.585 | 0.583 | 0.579 | 0.580 | 0.585 | 0.583 |
| DLC-ResNet50_s8 | 20 | b | 0.576 | 0.568 | 0.570 | 0.566 | 0.568 | 0.573 | 0.570 |
| DLC-ResNet50_s8 | 20 | d | 0.574 | 0.573 | 0.575 | 0.573 | 0.569 | 0.575 | 0.574 |
| DLC-EfficientNet-b7_s8 | 20 | c | **0.681** | **0.684** | **0.686** | **0.702** | **0.693** | **0.691** | **0.693** |
| DLC-EfficientNet-b7_s4 | 20 | d | 0.666 | 0.673 | 0.674 | 0.684 | 0.670 | 0.672 | 0.672 |
| DLC-EfficientNet-b7_s4 | 20 | c | 0.677 | 0.680 | 0.682 | 0.676 | 0.679 | 0.681 | 0.678 |

PW = paf width; G = Graph; c = calibrated; d = data-driven; b = baseline; s = stride; MF = multifusion; MS = multistage

**Table S5.**
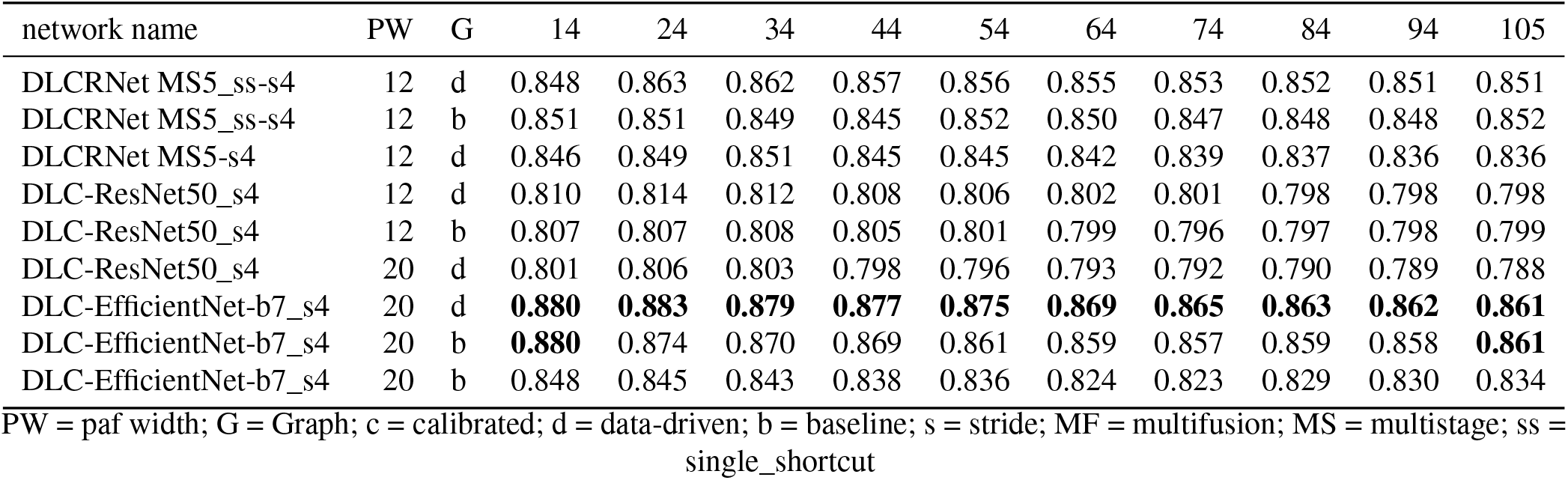
Mean Average Precision as a function of graph size for the marmoset dataset (more model benchmarking is available in the Suppl. Sheet).

| network name | PW | G | 14 | 24 | 34 | 44 | 54 | 64 | 74 | 84 | 94 | 105 |
| --- | --- | --- | --- | --- | --- | --- | --- | --- | --- | --- | --- | --- |
| DLCRNet MS5_ss-s4 | 12 | d | 0.848 | 0.863 | 0.862 | 0.857 | 0.856 | 0.855 | 0.853 | 0.852 | 0.851 | 0.851 |
| DLCRNet MS5_ss-s4 | 12 | b | 0.851 | 0.851 | 0.849 | 0.845 | 0.852 | 0.850 | 0.847 | 0.848 | 0.848 | 0.852 |
| DLCRNet MS5-s4 | 12 | d | 0.846 | 0.849 | 0.851 | 0.845 | 0.845 | 0.842 | 0.839 | 0.837 | 0.836 | 0.836 |
| DLC-ResNet50_s4 | 12 | d | 0.810 | 0.814 | 0.812 | 0.808 | 0.806 | 0.802 | 0.801 | 0.798 | 0.798 | 0.798 |
| DLC-ResNet50_s4 | 12 | b | 0.807 | 0.807 | 0.808 | 0.805 | 0.801 | 0.799 | 0.796 | 0.797 | 0.798 | 0.799 |
| DLC-ResNet50_s4 | 20 | d | 0.801 | 0.806 | 0.803 | 0.798 | 0.796 | 0.793 | 0.792 | 0.790 | 0.789 | 0.788 |
| DLC-EfficientNet-b7_s4 | 20 | d | **0.880** | **0.883** | **0.879** | **0.877** | **0.875** | **0.869** | **0.865** | **0.863** | **0.862** | **0.861** |
| DLC-EfficientNet-b7_s4 | 20 | b | **0.880** | 0.874 | 0.870 | 0.869 | 0.861 | 0.859 | 0.857 | 0.859 | 0.858 | **0.861** |
| DLC-EfficientNet-b7_s4 | 20 | b | 0.848 | 0.845 | 0.843 | 0.838 | 0.836 | 0.824 | 0.823 | 0.829 | 0.830 | 0.834 |

PW = paf width; G = Graph; c = calibrated; d = data-driven; b = baseline; s = stride; MF = multifusion; MS = multistage; ss = single\_shortcut

**Table S6.** Mean Average Precision as a function of graph size for the fish dataset (more model benchmarking is available in the Suppl. Sheet).

| network name | PW | G | 4 | 5 | 6 | 7 | 8 | 9 | 10 |
| --- | --- | --- | --- | --- | --- | --- | --- | --- | --- |
| DLCRNet MS5_ss-s4 | 12 | c | **0.683** | **0.692** | 0.681 | **0.677** | **0.685** | **0.660** | **0.654** |
| DLCRNet MS5_ss-s4 | 12 | d | 0.679 | 0.688 | 0.676 | 0.665 | 0.664 | 0.648 | 0.642 |
| DLCRNet MS5_shortcuts-s4 | 20 | d | 0.676 | 0.663 | 0.650 | 0.651 | 0.634 | 0.622 | 0.619 |
| DLC-ResNet50_s4 | 12 | d | 0.657 | 0.643 | 0.641 | 0.649 | 0.633 | 0.624 | 0.613 |
| DLC-ResNet50_s4 | 12 | c | 0.647 | 0.648 | 0.645 | 0.637 | 0.613 | 0.608 | 0.601 |
| DLC-ResNet50_s4 | 20 | c | 0.619 | 0.632 | 0.637 | 0.639 | 0.623 | 0.610 | 0.610 |
| DLC-EfficientNet-b0_s4 | 20 | d | 0.656 | 0.673 | **0.690** | 0.675 | 0.667 | 0.655 | 0.645 |
| DLC-EfficientNet-b0_s4 | 20 | c | 0.650 | 0.652 | 0.686 | 0.665 | 0.669 | 0.646 | 0.637 |
| DLC-EfficientNet-b0_s4 | 20 | b | 0.600 | 0.592 | 0.576 | 0.572 | 0.642 | 0.641 | 0.643 |

PW = paf width; G = Graph; c = calibrated; d = data-driven; b = baseline; s = stride; MF = multifusion; MS = multistage

**Figure S1.**
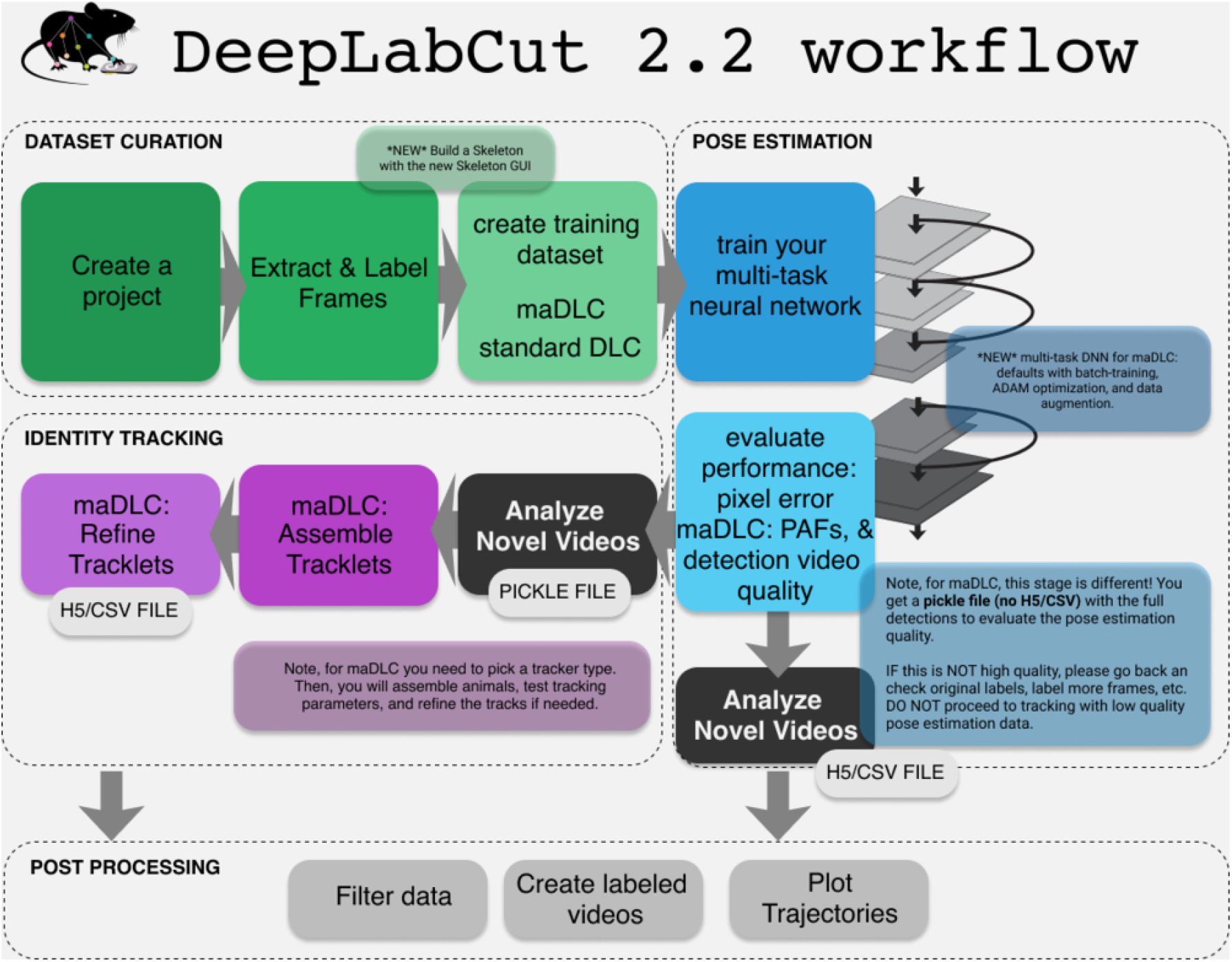
DeepLabCut 2.2 workflow. While the code is significantly updated, the user workflow is highly similar. Namely, we provide a full solution for dataset creation, labeling, train/test splits, neural network training, evaluation, and analysis (but now for multiple animals compared to previous versions). Here, we also provide new tools for all the 2.2. specific steps, such as creating tracklets, refining tracks, and output video creation tools.

**Figure S2.**
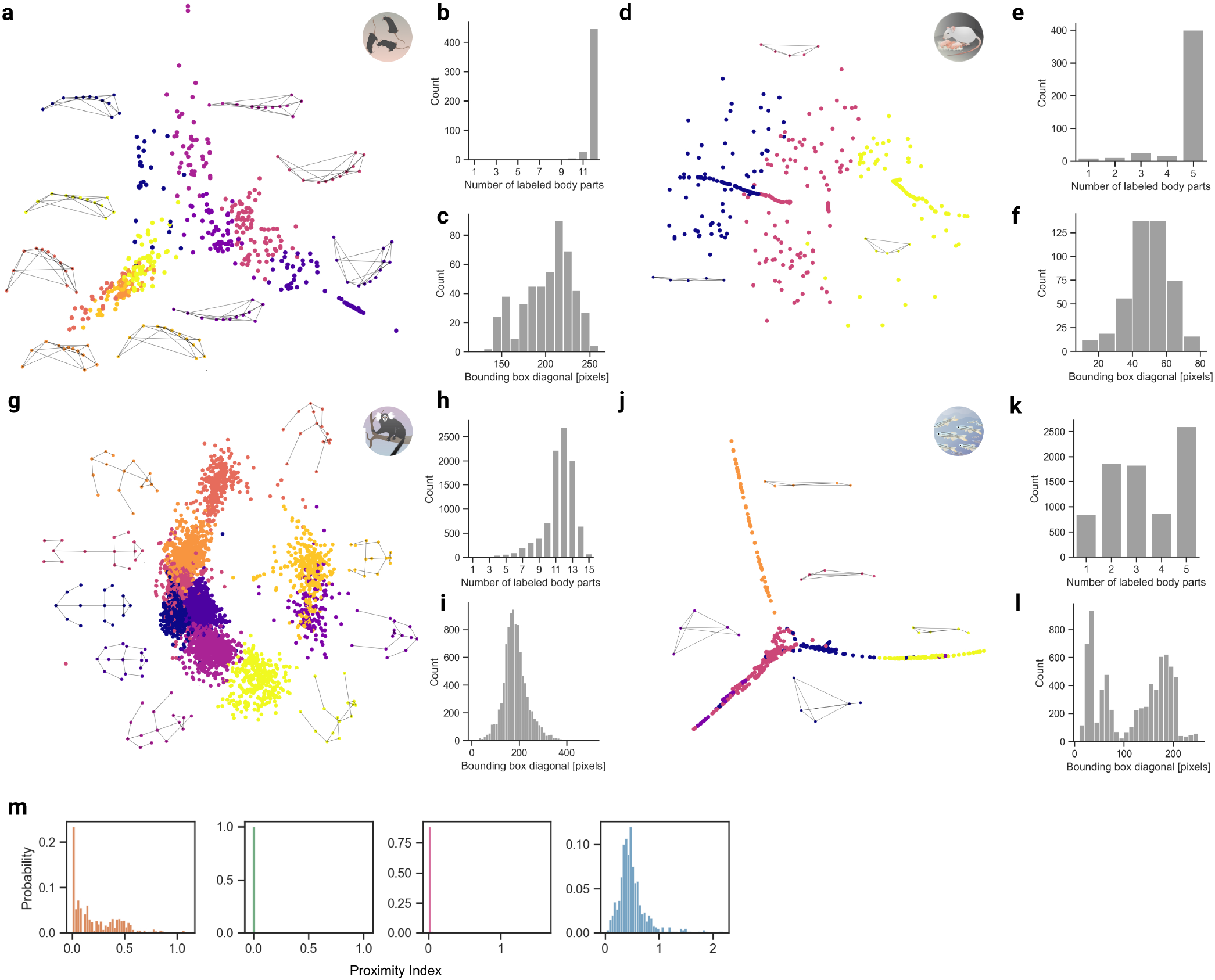
Datasets characteristics and statistics for four datasets. Normalized animal poses were clustered using K-means adapted for missing elements, and embedded non-linearly in 2D space via Isometric mapping (54); representative poses are also shown **(a,d,g,j)**. Counts of labeled keypoints **(b,e,h,k)** and distribution of bounding box diagonal lengths **(c,f,i,l)**. Remarkably, the range of the marmoset scale (50 400 pixels) is much wider than all the other datasets. Note the bimodality of the fish diagonals **(l)**, characteristic of side-to-side flapping and marked changes in body conformation. The Proximity Index (m) reflects the crowdedness of the various dataset scenes, with the mice and fish being more cluttered on average than the pups and marmosets. Statistics were computed from the ground truth test video annotations.

**Figure S3.**
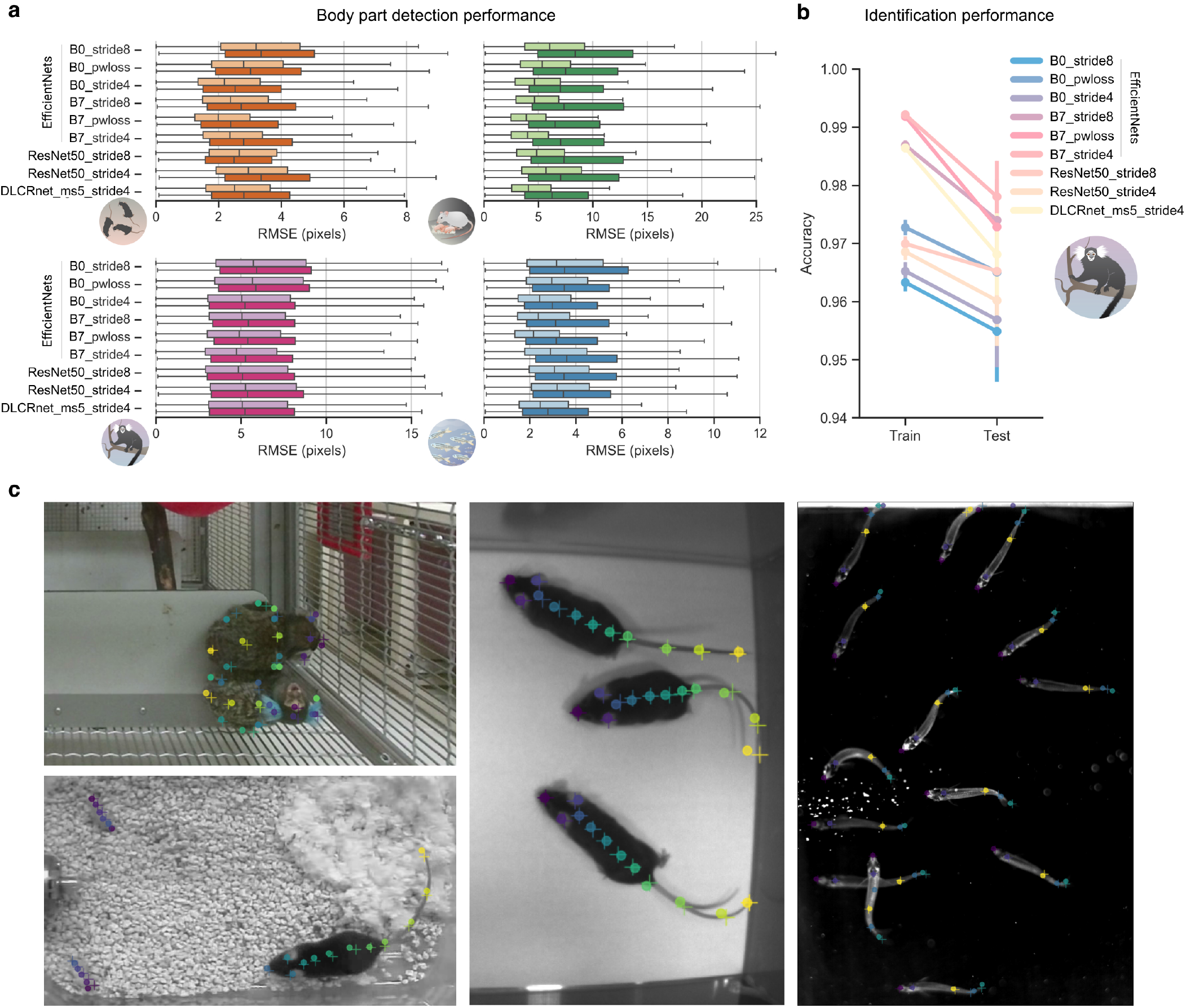
Performance of various DeepLabCut network architectures. **(a)**: Overall keypoint prediction errors of ResNets-50 and the EfficientNets backbones (B0/B7), at stride 4 and 8. Distribution of train and test errors are displayed as light and dark boxplots, respectively. **(b)**: Marmoset identification train–test accuracy *vs* backbones. **(c)**: Images on held-out test data, where “+” is human ground truth, and the circle is model prediction (shown for ResNet50_stride 8).

**Figure S4.**
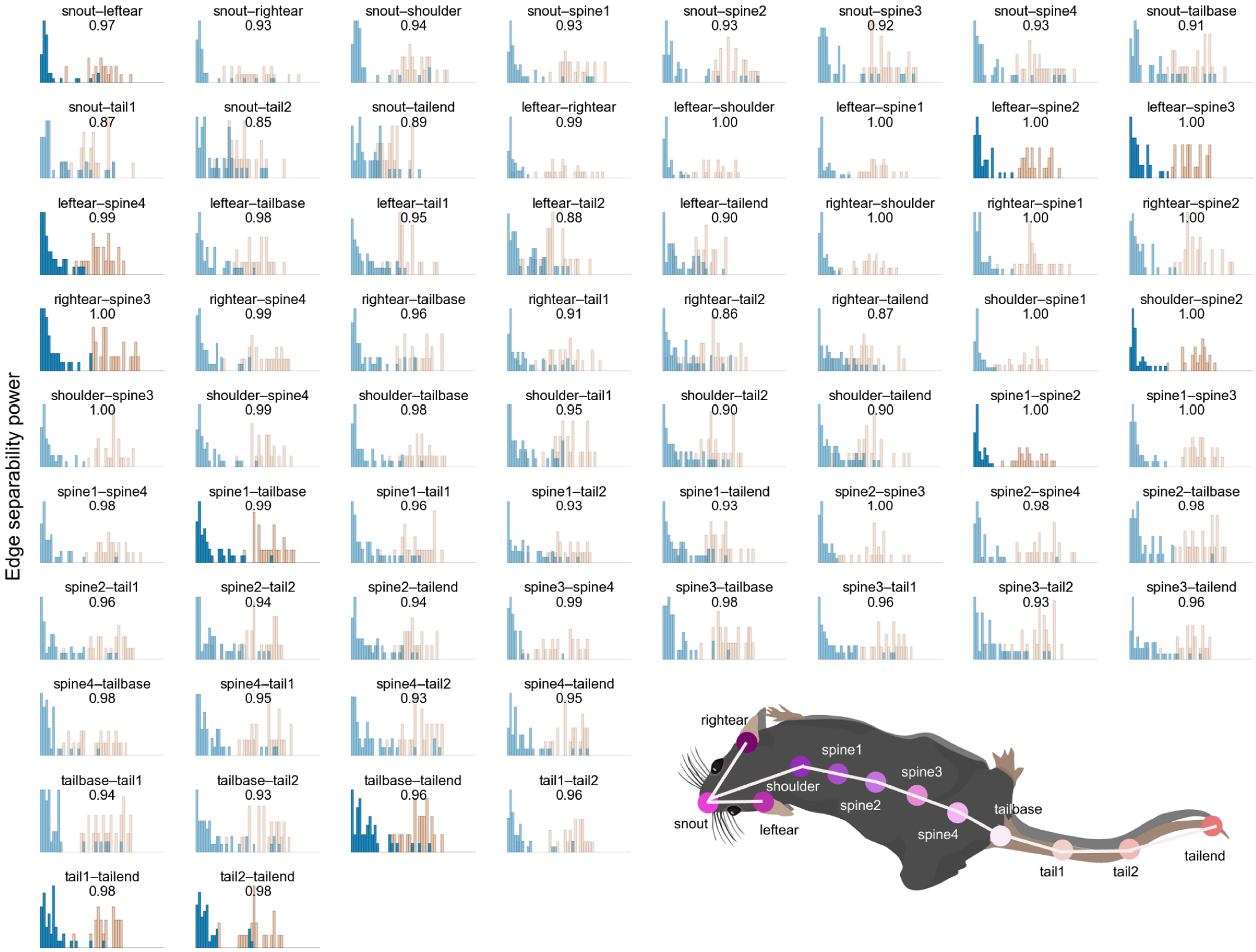
Discriminability of part affinity fields. Within- (pink) and between-animal (blue) affinity cost distributions for all edges of the mouse skeleton with ResNet50_stride8. The saturated subplots highlight the 11 edges kept to form the smallest, optimal part affinity graph (see Figure 2b). This is based on the separability power of an edge, i.e., its ability to discriminate a connection between two keypoints effectively belonging to the same animal from the wrong ones, and reflected by the corresponding AUC scores (at the top of the subplots).

**Figure S5.**
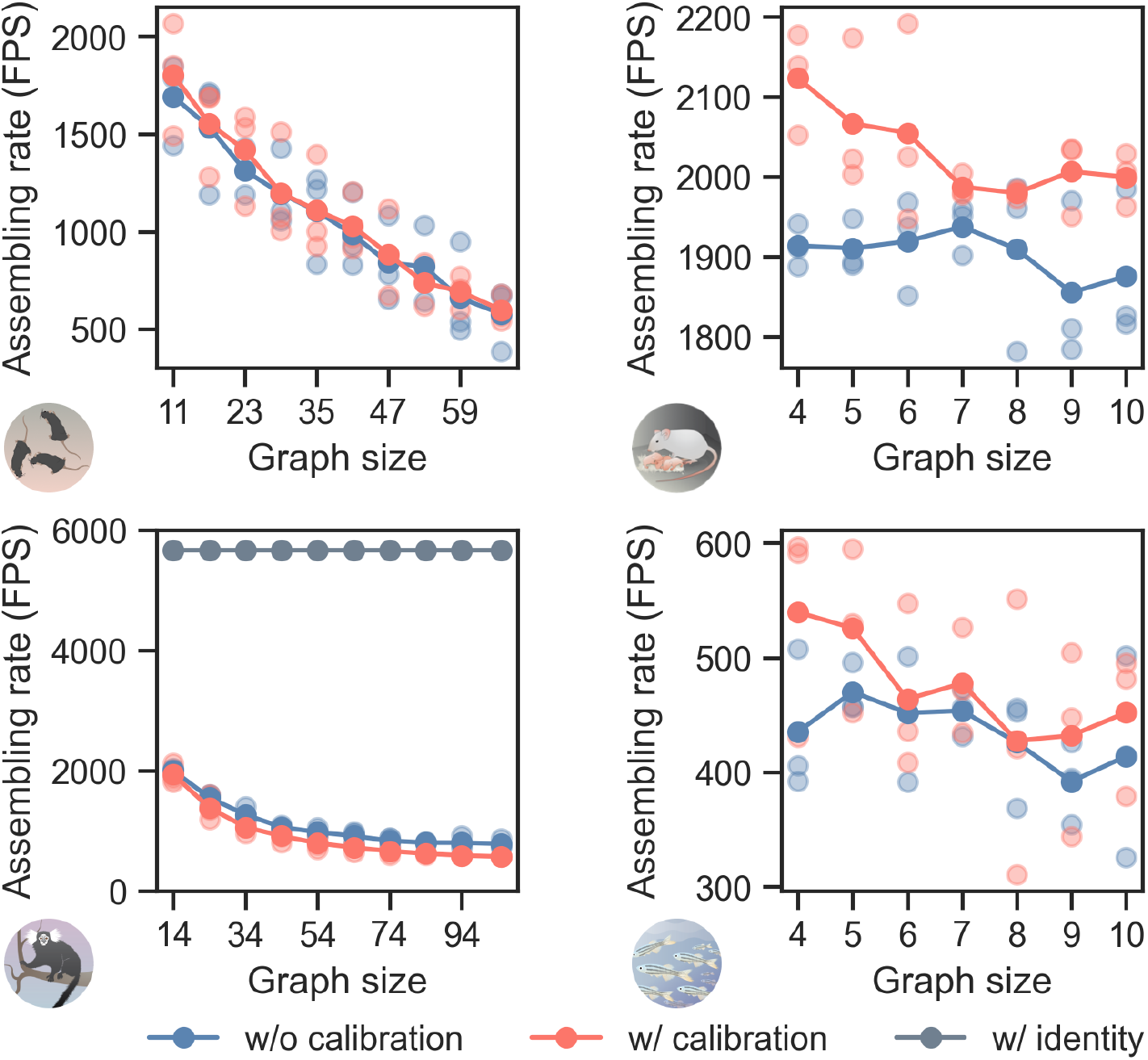
Average animal assembly speed in frames per second as a function of graph size. Speed decays approximately with the cube of the graph size. Improving the assembly robustness via calibration with labeled data in large graphs incurs no extra computational cost at best, and a slowdown by 25% at worst; remarkably, it is found to accelerate assembly speed in small graphs. Relying exclusively on keypoint identity prediction results in average speeds of ~ 5600 frames per second, independent of graph size. Three timing experiments were run per graph size (lighter colored dots). Note that assembling rates exclude CNN processing times. Speed benchmark was run on a workstation with an Intel(R) Core(TM) i9-10900X CPU @ 3.70GHz.

**Figure S6.**
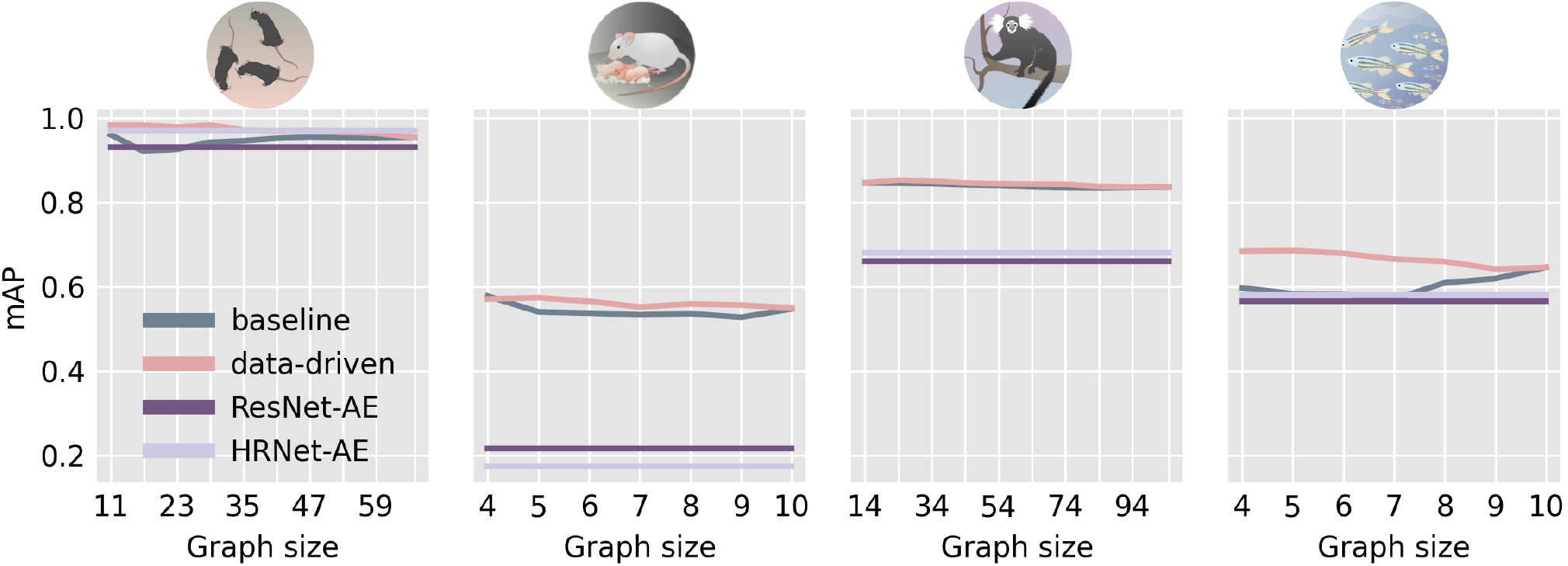
Comparison to state-of-the-art methods on COCO. Mean Average Precision (mAP) as a function of graph size; note that the associative embedding method does not rely on a graph (and thus graph size). The performance of MMPose’s implementation of ResNet-AE and HRNet-AE bottom-up variants is shown for comparison against our multi-stage architecture (DLCRNet_ms5). Only for the tri-mouse dataset do these state-of-the-art networks with associative embedding perform on par with our data-driven assembling method and only with the HRNet backbone.

## Footnotes

1 For example, 10 keypoints yield 261,080 different possible connected graphs http://oeis.org/A001349; admittedly all are not suitable for pose estimation.

## References

1. Roland Kays, Margaret C Crofoot, Walter Jetz, and Martin Wikelski. Terrestrial animal tracking as an eye on life and planet. Science, 348(6240):aaa2478, 2015.

2. Daniel Schofield, Arsha Nagrani, Andrew Zisserman, Misato Hayashi, Tetsuro Matsuzawa, Dora Biro, and Susana Carvalho. Chimpanzee face recognition from videos in the wild using deep learning. Science advances, 5(9):eaaw0736, 2019.

3. Mohammad Sadegh Norouzzadeh, Anh Nguyen, Margaret Kosmala, Alexandra Swanson, Meredith S Palmer, Craig Packer, and Jeff Clune. Automatically identifying, counting, and describing wild animals in camera-trap images with deep learning. Proceedings of the National Academy of Sciences, 115(25):E5716–E5725, 2018.

4. Maxime Vidal, Nathan Wolf, Beth Rosenberg, Bradley P Harris, and Alexander Mathis. Perspectives on individual animal identification from biology and computer vision. arXiv preprint arXiv:2103.00560, 2021.

5. Sandeep Robert Datta, David J Anderson, Kristin Branson, Pietro Perona, and Andrew Leifer. Computational neuroethology: a call to action. Neuron, 104(1):11–24, 2019.

6. Mackenzie Weygandt Mathis and Alexander Mathis. Deep learning tools for the measurement of animal behavior in neuroscience. Current Opinion in Neurobiology, 60:1–11, 2020.

7. Zhe Cao, Tomas Simon, Shih-En Wei, and Yaser Sheikh. Realtime multi-person 2d pose estimation using part affinity fields. In Proceedings of the IEEE conference on computer vision and pattern recognition, 2017.

8. Alejandro Newell, Zhiao Huang, and Jia Deng. Associative embedding: End-to-end learning for joint detection and grouping. In Advances in Neural Information Processing Systems, pages 2277–2287, 2017.

9. Bowen Cheng, Bin Xiao, Jingdong Wang, Honghui Shi, Thomas S Huang, and Lei Zhang. Higherhrnet: Scale-aware representation learning for bottom-up human pose estimation. In Proceedings of the IEEE/CVF Conference on Computer Vision and Pattern Recognition, pages 5386–5395, 2020.

10. Lucas Stoffl, Maxime Vidal, and Alexander Mathis. End-to-end trainable multi-instance pose estimation with transformers. arXiv preprint arXiv:2103.12115, 2021.

11. Eldar Insafutdinov, Leonid Pishchulin, Bjoern Andres, Mykhaylo Andriluka, and Bernt Schiele. DeeperCut: A deeper, stronger, and faster multi-person pose estimation model. In European Conference on Computer Vision, pages 34–50. Springer, 2016.

12. Sven Kreiss, Lorenzo Bertoni, and Alexandre Alahi. Pifpaf: Composite fields for human pose estimation. In Proceedings of the IEEE Conference on Computer Vision and Pattern Recognition, pages 11977–11986, 2019.

13. Jiefeng Li, Can Wang, Hao Zhu, Yihuan Mao, Hao-Shu Fang, and Cewu Lu. Crowdpose: Efficient crowded scenes pose estimation and a new benchmark. In Proceedings of the IEEE Conference on Computer Vision and Pattern Recognition, pages 10863–10872, 2019.

14. Jingdong Wang, Ke Sun, Tianheng Cheng, Borui Jiang, Chaorui Deng, Yang Zhao, Dong Liu, Yadong Mu, Mingkui Tan, Xinggang Wang, et al. Deep high-resolution representation learning for visual recognition. IEEE transactions on pattern analysis and machine intelligence, 2020.

15. Alexander Mathis, Steffen Schneider, Jessy Lauer, and Mackenzie W. Mathis. A primer on motion capture with deep learning: Principles, pitfalls, and perspectives. Neuron, 108:44–65, 2020.

16. Cristina Segalin, Jalani Williams, Tomomi Karigo, May Hui, Moriel Zelikowsky, Jennifer J Sun, Pietro Perona, David J Anderson, and Ann Kennedy. The mouse action recognition system (mars): a software pipeline for automated analysis of social behaviors in mice. bioRxiv, 2020.

17. Talmo D Pereira, Nathaniel Tabris, Junyu Li, Shruthi Ravindranath, Eleni S Papadoyannis, Z Yan Wang, David M Turner, Grace McKenzie-Smith, Sarah D Kocher, Annegret Lea Falkner, et al. Sleap: multi-animal pose tracking. bioRxiv, 2020.

18. Zexin Chen, Ruihan Zhang, Yu Eva Zhang, Haowen Zhou, Hao-Shu Fang, Rachel R Rock, Aneesh Bal, Nancy Padilla-Coreano, Laurel Keyes, Kay M Tye, et al. Alphatracker: A multi-animal tracking and behavioral analysis tool. bioRxiv, 2020.

19. Tsung-Yi Lin, Michael Maire, Serge Belongie, James Hays, Pietro Perona, Deva Ramanan, Piotr Dollár, and C Lawrence Zitnick. Microsoft coco: Common objects in context. In European conference on computer vision, pages 740–755. Springer, 2014.

20. Alexander Mathis, Pranav Mamidanna, Kevin M Cury, Taiga Abe, Venkatesh N Murthy, Mackenzie Weygandt Mathis, and Matthias Bethge. Deeplabcut: markerless pose estimation of user-defined body parts with deep learning. Nature neuroscience, 21:1281–1289, 2018.

21. Tanmay Nath, Alexander Mathis, An Chi Chen, Amir Patel, Matthias Bethge, and Mackenzie W Mathis. Using deeplabcut for 3d markerless pose estimation across species and behaviors. Nature protocols, 14:2152–2176, 2019.

22. Kaiming He, Xiangyu Zhang, Shaoqing Ren, and Jian Sun. Deep residual learning for image recognition. In Proceedings of the IEEE conference on computer vision and pattern recognition, pages 770–778, 2016.

23. Mingxing Tan and Quoc Le. Efficientnet: Rethinking model scaling for convolutional neural networks. In International Conference on Machine Learning, pages 6105–6114, 2019.

24. Kunal K Ghosh, Laurie D Burns, Eric D Cocker, Axel Nimmerjahn, Yaniv Ziv, Abbas El Gamal, and Mark J Schnitzer. Miniaturized integration of a fluorescence microscope. Nature methods, 8(10):871, 2011.

25. Alex Bewley, Zongyuan Ge, Lionel Ott, Fabio Ramos, and Ben Upcroft. Simple online and realtime tracking. In 2016 IEEE International Conference on Image Processing (ICIP), pages 3464–3468. IEEE, 2016.

26. M Bertozzi, A Broggi, A Fascioli, A Tibaldi, R Chapuis, and F Chausse. Pedestrian localization and tracking system with kalman filtering. In IEEE Intelligent Vehicles Symposium, 2004, pages 584–589. IEEE, 2004.

27. Gary A Kane, Gonçalo Lopes, Jonny L Saunders, Alexander Mathis, and Mackenzie W Mathis. Real-time, low-latency closed-loop feedback using markerless posture tracking. Elife, 9:e61909, 2020.

28. Alexander Mathis, Thomas Biasi, Steffen Schneider, Mert Yuksekgonul, Byron Rogers, Matthias Bethge, and Mackenzie W Mathis. Pretraining boosts out-of-domain robustness for pose estimation. In Proceedings of the IEEE/CVF Winter Conference on Applications of Computer Vision, pages 1859–1868, 2021.

29. Gines Hidalgo, Yaadhav Raaj, Haroon Idrees, Donglai Xiang, Hanbyul Joo, Tomas Simon, and Yaser Sheikh. Single-network wholebody pose estimation. arXiv preprint arXiv:1909.13423, 2019.

30. Francisco Romero-Ferrero, Mattia G Bergomi, Robert C Hinz, Francisco JH Heras, and Gonzalo G de Polavieja. idtracker. ai: tracking all individuals in small or large collectives of unmarked animals. Nature methods, 16(2):179, 2019.

31. Tristan Walter and Ian D Couzin. Trex, a fast multi-animal tracking system with markerless identification, and 2d estimation of posture and visual fields. Elife, 10:e64000, 2021.

32. Xiongwei Wu, Doyen Sahoo, and Steven CH Hoi. Recent advances in deep learning for object detection. Neurocomputing, 2020.

33. Jacob M Graving, Daniel Chae, Hemal Naik, Liang Li, Benjamin Koger, Blair R Costelloe, and Iain D Couzin. Deepposekit, a software toolkit for fast and robust animal pose estimation using deep learning. eLife, 8:e47994, oct 2019. ISSN 2050-084X. doi: 10.7554/eLife.47994.

34. Zheng Wu, Anita E Autry, Joseph F Bergan, Mitsuko Watabe-Uchida, and Catherine G Dulac. Galanin neurons in the medial preoptic area govern parental behaviour. Nature, 509(7500):325, 2014.

35. Johannes Kohl, Benedicte M Babayan, Nimrod D Rubinstein, Anita E Autry, Brenda Marin-Rodriguez, Vikrant Kapoor, Kazunari Miyamishi, Larry S Zweifel, Liqun Luo, Naoshige Uchida, et al. Functional circuit architecture underlying parental behaviour. Nature, 556(7701):326, 2018.

36. Valentina Di Santo, Erin L Blevins, and George V Lauder. Batoid locomotion: effects of speed on pectoral fin deformation in the little skate, *leucoraja erinacea*. Journal of Experimental Biology, 220(4): 705–712, 2017.

37. Mark Sandler, Andrew Howard, Menglong Zhu, Andrey Zhmoginov, and Liang-Chieh Chen. Mobilenetv2: Inverted residuals and linear bottlenecks. In Proceedings of the IEEE Conference on Computer Vision and Pattern Recognition, pages 4510–4520, 2018.

38. Diederik P Kingma and Jimmy Ba. Adam: A method for stochastic optimization. arXiv preprint arXiv:1412.6980, 2014.

39. Eldar Insafutdinov, Mykhaylo Andriluka, Leonid Pishchulin, Siyu Tang, Evgeny Levinkov, Bjoern Andres, and Bernt Schiele. Arttrack: Articulated multi-person tracking in the wild. In Proceedings of the IEEE conference on computer vision and pattern recognition, 2017.

40. Benjamin Biggs, Thomas Roddick, Andrew Fitzgibbon, and Roberto Cipolla. Creatures great and smal: Recovering the shape and motion of animals from video. In Asian Conference on Computer Vision, pages 3–19. Springer, 2018.

41. Yi Yang and Deva Ramanan. Articulated human detection with flexible mixtures of parts. IEEE transactions on pattern analysis and machine intelligence, 35(12):2878–2890, 2012.

42. Anna Huang. Similarity measures for text document clustering. In Proceedings of the sixth new zealand computer science research student conference (NZCSRSC2008), Christchurch, New Zealand, volume 4, pages 9–56, 2008.

43. Raphael Vallat. Pingouin: statistics in python. Journal of Open Source Software, 3(31):1026, 2018.

44. Ke Sun, Bin Xiao, Dong Liu, and Jingdong Wang. Deep high-resolution representation learning for human pose estimation. arXiv preprint arXiv:1902.09212, 2019.

45. MMPose Contributors. Openmmlab pose estimation toolbox and benchmark.https://github.com/open-mmlab/mmpose, 2020.

46. Rohit Girdhar, Georgia Gkioxari, Lorenzo Torresani, Manohar Paluri, and Du Tran. Detect-and-track: Efficient pose estimation in videos. In Proceedings of the IEEE Conference on Computer Vision and Pattern Recognition, pages 350–359, 2018.

47. Patrick Emami, Panos M Pardalos, Lily Elefteriadou, and Sanjay Ranka. Machine learning methods for solving assignment problems in multi-target tracking. arXiv preprint arXiv:1802.06897, 2018.

48. Li Zhang, Yuan Li, and Ramakant Nevatia. Global data association for multi-object tracking using network flows. In 2008 IEEE Conference on Computer Vision and Pattern Recognition, pages 1–8. IEEE, 2008.

49. Aric A. Hagberg, Daniel A. Schult, and Pieter J. Swart. Exploring network structure, dynamics, and function using networkx. In Gaël Varoquaux, Travis Vaught, and Jarrod Millman, editors, Proceedings of the 7th Python in Science Conference, pages 11 – 15, Pasadena, CA USA, 2008.

50. Caglayan Dicle, Octavia I Camps, and Mario Sznaier. The way they move: Tracking multiple targets with similar appearance. In Proceedings of the IEEE international conference on computer vision, pages 2304–2311, 2013.

51. Huanhuan Yin, Zhifang Zhu, and Feng Ding. Model order determination using the hankel matrix of impulse responses. Applied Mathematics Letters, 24(5):797–802, 2011.

52. Ravindra K Ahuja, Thomas L Magnanti, and James B Orlin. Network flows: theory, algorithms, and applications. Prentice-Hall, 1993.

53. Keni Bernardin and Rainer Stiefelhagen. Evaluating multiple object tracking performance: the clear mot metrics. EURASIP Journal on Image and Video Processing, 2008:1–10, 2008.

54. Joshua B Tenenbaum, Vin De Silva, and John C Langford. A global geometric framework for nonlinear dimensionality reduction. science, 290(5500):2319–2323, 2000.

